# RAS pathway activation drives clonal selection and monocytic differentiation in FLT3 and BCL2 inhibitor resistance

**DOI:** 10.1101/2025.02.02.636108

**Authors:** Vanessa E. Kennedy, Cheryl A. C. Peretz, Anushka Walia, Brenda Chyla, Yan Sun, Jason Hill, Elaine Tran, Andrew Koh, Timothy Ferng, Samantha Pintar, Matthew Jones, Bogdan Popescu, Natalia Murad, Ritu Roy, Adam Olshen, Sunil Joshi, Elie Traer, Monique Dail, Habib Hamidi, Jessica Altman, Naval Daver, Mark Levis, James McCloskey, Alexander Perl, Catherine C. Smith

**Affiliations:** Stanford University; university of California San Francisco; AbbVie; Astellas Pharma; Oregon Health & Science University; Genentech, Inc.; Northwestern University; University of Texas MD Anderson Cancer Center; Johns Hopkins University; John Theurer Cancer Center at Hackensack University Medical Center; Perelman School of Medicine and the Abramson Cancer Center at the University of Pennsylvania

## Abstract

Despite efficacy of FLT3 and BCL2 inhibition in acute myeloid leukemia (AML), relapse limits survival. Mutation status and AML monocytic differentiation are implicated in resistance. On-treatment tumor evolution may select for genetically distinct clones or shifts in differentiation not resolvable by bulk sequencing. We performed multiomic single cell (SC) DNA/protein and RNA/protein profiling of patients treated on a clinical trial of the BCL2 inhibitor venetoclax and the FLT3 inhibitor gilteritinib (Ven/Git) to characterize immunophenotypic, transcriptional, and genetic clonal evolution on therapy. We found that while Ven/Gilt effectively eliminated FLT3 mutant clones, it selected for RAS mutations, RAS pathway activation and RAS-associated monocytic differentiation. In an *in vitro* model of monocytic differentiation associated with heightened RAS pathway activation, we demonstrated that MEK inhibition re-sensitized to Ven/Gilt. These data indicate RAS signaling is central to FLT3 and BCL2 inhibitor resistance, is tightly coupled to monocytic differentiation and can be overcome by RAS pathway inhibition.

**COI:** C.C.S. has provided educational talks for Astellas Pharma, served on advisory boards for Genentech/Abbvie and received research funding from Abbvie. B.C., Y.S. and J.H. are employees of Abbvie. M.S. and H.H. are or were previously employees of Genentech.

**Funding:** This work was supported in part by Abbvie. C.C.S. is a Leukemia & Lymphoma Society Scholar in Clinical Research and a Damon Runyon-Richard Lumsden Foundation Clinical Investigator supported (in part) by the Damon Runyon Cancer Research Foundation (CI-99–18).

**Statement of Significance:** Mutational and non-mutational RAS signaling activation drives clonal selection, monocytic differentiation and treatment resistance to FLT3 and BCL2 inhibition in AML. MEK inhibition can resensitize resistant AML cells, suggesting therapeutic potential for combined FLT3, BCL2 and RAS pathway inhibition in AML.

## Introduction

Acute myeloid leukemia (AML) is a devastating hematologic malignancy driven by heterogeneous molecular mechanisms. AML treatment has traditionally consisted of cytotoxic chemotherapy; however, as specific biologic drivers have emerged, treatment has evolved to incorporate targeted therapies. These include inhibitors targeted to specific pathogenic mutations, such as *FLT3*, *IDH1*, or *IHD2*, as well as the BCL2 inhibitor venetoclax, which has efficacy across broad genetic subtypes. These targeted therapies, either alone or in combination, have transformed the treatment landscape of both newly diagnosed and relapsed/refractory (R/R) AML^1–6^. While promising, targeted therapies are not curative, and relapse remains common. Understanding the mechanisms of resistance to targeted therapies is critical to improve patient outcomes.

For patients with AML, *FLT3* mutations have historically conferred a poor prognosis, principally due to an increased risk of relapse^7^. The FLT3 tyrosine kinase inhibitor gilteritinib is now standard of care for R/R FLT3-mutant AML^8^, but resistance develops in most patients. Recently, the addition of venetoclax to gilteritinib was shown to nearly double response rates compared to single agent gilteritinib^9^. However, even with venetoclax/gilteritinib (Ven/Gilt) combination, patients eventually relapse absent stem cell transplantation^9^. Development of therapeutic strategies for AML resistant to Ven/Gilt and other targeted therapy combinations requires thorough understanding of resistance mechanisms.

Although the mechanisms of resistance to the Ven/Gilt combination have not been elucidated, resistance to each drug has been independently described. Signaling mutations, particularly RAS pathway mutations, have been implicated in resistance to multiple targeted therapies, including gilteritinib^10^ and venetoclax^11,12^. However, mutations account for only a subset of resistance^13,14^. Further, there is increasing awareness that cellular differentiation state may impact sensitivity to AML targeted therapy. While in some therapies stemness connotes resistance^15^, monocytic differentiation has been linked to intrinsic resistance to venetoclax^16,17^ and to FLT3 inhibitors *in vitro*^18^. Monocytic differentiation has also previously been correlated with RAS mutation^19^ and activation^20^, but this correlation is imperfect. It is also unknown if monocytic differentiation alone is sufficient to drive resistance to either BCL2 or FLT3 inhibition. Multiple groups have described a variety of potential mechanisms underlying the role of AML monocytic differentiation in venetoclax resistance, including increased expression pro-apoptotic proteins such as MCL1 and BCL2A1 and increased reliance on oxidative phosphorylation or purine metabolism^17^. However, the full molecular mechanisms by which monocytic differentiation mediates resistance to venetoclax and how these might translate to resistance to other therapies remains uncertain. Further, while both genetic drivers and cellular differentiation state may influence resistance to targeted therapies, how these factors interact remains unknown. More importantly, therapeutic approaches to overcome these resistance mechanisms remain unproven.

Single cell (SC) DNA sequencing (DNAseq) has shown that resistance to targeted therapy can involve selection for heterogeneous mutant resistant AML clones, some of which are pre-existing^10,11,21,22^. However, despite emerging data that AML differentiation state can impact sensitivity to therapy^17^, dynamic changes in the differentiation state of AML clones after targeted therapy have not been documented. We used multiomic SC DNA, RNA, and immunophenotypic profiling of longitudinal samples from patients treated on a phase 1b trial of Ven/Gilt (NCT03625505) to characterize immunophenotypic, transcriptional, and genetic clonal evolution. We found that while Ven/Gilt effectively reduced or eliminated FLT3 mutant clones, including those with other oncogenic driver co-mutations (i.e. IDH1/2 mutations), the combination strongly selected for RAS mutations as well as non-mutational RAS pathway activation and RAS-associated monocytic differentiation. These findings implicate a central role for RAS signaling activation in resistance to both FLT3 and BCL2 inhibitor therapy that is tightly coupled to monocytic differentiation programs. In an *in vitro* model of increased monocytic differentiation associated with heightened RAS pathway activation, we further demonstrated the ability of MEK inhibition to re-sensitize to Ven/Gilt. These data strongly suggest that therapies effectively targeting RAS signaling are required to overcome resistance to both FLT3 and BCL2 targeted inhibitors, regardless of RAS mutation status.

## Results

### The genetic landscape of R/R FLT3-mutant AML

We profiled 35 pre-Ven/Gilt, on-treatment, and, if available, relapse samples from 12 patients with R/R, *FLT3-*mutated AML (2-5 samples/patient) (**Figure 1A).** All patients were enrolled on the phase Ib/II trial of gilteritinib plus venetoclax (NCT03625505) (**Table S1, Figure S1)**^9^. Of the 12 patients, 8 had received prior targeted therapy with a tyrosine kinase inhibitor (TKI) and 1 had received prior venetoclax. Nine patients had clinical response to Ven/Gilt with a median relapse-free time of 8.3 months; 7 of these patients ultimately developed evidence of relapse (**Figure 1B**). We profiled all 35 samples via SC DNAseq, for a total of 107,528 single cells (median 8,960 cells/patient); 18 samples were jointly profiled via SC DNAseq and cell-surface immunophenotyping (SC DAbseq). We used a custom panel targeting hotspots in 66 genes frequently mutated in AML or attributed to targeted therapy resistance in other cancers in conjunction with 9 antibody-oligonucleotide conjugates for cell-surface immunophenotypic proteins on hematopoietic cells (**Table S2, S3**).

**Figure 1.**
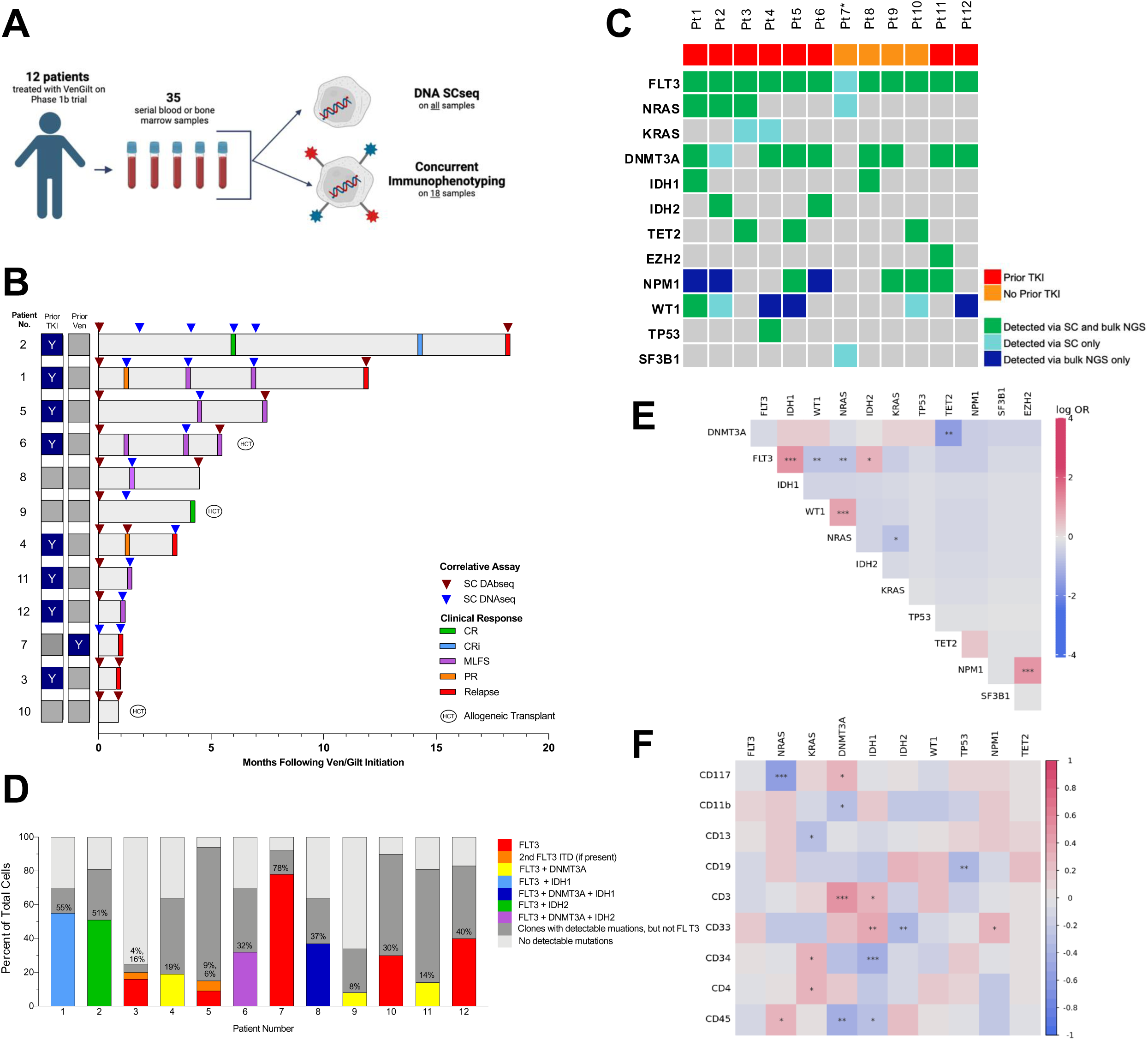
Clinical and Genetic Landscape of 12 patients with relapsed/refractory (R/R) acute myeloid leukemia (AML) treated with Venetoclax and Gilteritinib (Ven/Gilt). **A.** Schematic depicting sample workflow for single-cell Dabseq. Ven/Gilt, venetoclax and gilteritinib; SCseq, single-cell sequencing **B.** Swimmer’s plot of 12 patient cohort treated on clinical trial. Each horizontal bar is a unique patient. Clinical responses were evaluated on the basis of guidelines adapted from the International Working Group for AML^83^. Triangles indicate correlative samples. Bars to the left of the Y axis indicate treatment with TKI and/or venetoclax prior to trial enrollment. TKI, tyrosine kinase inhibitor; Ven, venetoclax; CR, complete response; CRi, complete response with incomplete hematologic recovery; MLFS, morphological leukemia-free state; PR, progressive disease. **C.** Oncoplot of the 12 patient cohort prior to Ven/Gilt therapy. Each column is a unique patient. Mutations are color-coded based on whether they were detected via bulk next-generation sequencing, single-cell DNA sequencing, or both. Only mutations covered by the SC amplicon panel are depicted. *Patient 7 did not have bulk sequencing available. **D.** Bar chart depicting diversity of clonal composition and size of *FLT3-*mutated clones prior to Ven/Gilt. Bars are color-coded based on *FLT3* mutation(s) and co-mutations, and percentages reflect relative proportion of *FLT3-*mutated clones for individual patients. Dark gray sections represent clones with mutations detected aside from *FLT3* and light gray sections represent sequenced cells without pathogenic mutations detected by the SC sequencing panel. **E.** Pairwise association of driver mutations identified via SC DNA sequencing across 48 clones in 12 patient cohort prior to Ven/Gilt therapy. For each mutation pair, cooccurrence is summarized as log odds ratio (OR), with positive values indicating cooccurrence and negative values mutual exclusivity. Statistical significance is indicated as *p, .05; **p, .01; ***p, .001. **F.** Correlation matrix of association between mutations and 9 cell-surface proteins in 11 patients prior to Ven/Gilt. The plot shows cooccurence (red) or exclusivity (blue) based on point-biserial estimate with statistician significance indicated as *p , .05; **p , .01; ***p , .001.

The baseline mutational landscape for all patients is depicted in **Figure 1C-F**. All patients had mutations in *FLT3* as per trial eligibility; other common pre-treatment mutations included mutations in *DNMT3A* (9 patients), *NPM1* (7), *WT1* (6), and *NRAS* (4) (**Figure 1C**). At baseline and prior to Ven/Gilt therapy, we identified 48 individual genetically defined subclones, for a median of 4.5 clones per patient (range 2 - 5). Although all patients harbored *FLT3* mutations, only 29% (14/48) of clones contained *FLT3* mutations.

Across all 12 patients, we detected 14 distinct *FLT3-*mutated clones; 2 patients harbored 2 distinct *FLT3-ITD-*mut populations that were not co-mutated in the same single cell or clone (**Figure 1D**). These 14 *FLT3-*mut clones demonstrated diverse pre-treatment sizes, ranging from 4 - 78% of all sequenced cells, and represented the major clone in some patients (e.g., Patient 7) and the minor clone in others (e.g. Patient 10). Within the *FLT3-*mutated clones, the most common co-mutated genes were *DNMT3A* (5/14 clones) and *IDH1/2* (each 2/14 clones). Across all 48 identified clones, clone-level mutational co-occurrence analysis demonstrated the strongest positive association between *FLT3* and *IDH1* (Odds Ratio [OR] 12.60, p = <0.001), *NRAS*/*WT1* (OR = 7.94, p = <0.001), and *NPM1*/*EZH2* (OR = 17.8, p = <0.001); negative associations were observed between *TP53/TET2* (OR = 0.02, p = 0.008), *FLT3/NRAS* (OR = 0.25, p = 0.009), *FLT3/WT1* and (OR = 0.28, p = 0.012) (**Figure 1E**).

### Heterogenous genotype-immunophenotype associations in R/R FLT3-mutated AML

Using SC DAbseq, we next investigated the associations between genotype and immunophenotype in R/R *FLT3-*mutated AML prior to Ven/Gilt therapy. Across the 11 patients and 45 clones with mult-iomic profiling available at baseline, we observed several broad genotype-immunophenotype associations (**Figure 1F**). At the clonal level, we observed considerable inter- and intra-patient heterogeneity (**Figure S2**). For example, Patient 3 harbored 2 distinct *FLT3-ITD*-mutated subclones without detected co-mutations. Despite having similar driver mutations, the two *FLT3-ITD*-mutated subclones demonstrated significantly different immunophenotypes; one *FLT3-ITD-*mutated clone had an immature immunophenotype with high CD34 and CD117 expression, while the second demonstrated a more monocytic immunophenotype with higher expression of CD13 and CD11b (**Figure S2 A - C**). In Patient 2, however, genotype and immunophenotype were not associated. Despite having distinct CD117+/CD34+ and CD13+/CD33+ subpopulations, these populations were not significantly enriched with any specific driver mutations, although it is possible mutations were present in genes not covered by our panel (**Figure S2 D - F)**.

### Clonal evolution of genotype with Ven/Gilt eradicates FLT3-mutant clones, including in the presence of co-mutations

Using longitudinal SC DNAseq, we next sought to profile evolution of genetic clonal architecture on Ven/Gilt treatment. Across all patients and all timepoints, we identified 50 genetically distinct subclones (**Figure 2A; Figure S3; Table S4**). Ven/Gilt was active against *FLT3-*mutated clones: all 14 *FLT3-*mutated clones decreased in size on therapy, and at time of maximal response, 11 clones from 9 patients decreased to an undetectable level (**Figure 2B**). Responses of the *FLT3*-mutated clones were rapid and frequently decreased by cycle 1 day 28 in 8 patients. Only one *FLT3-*mutated clone subsequently recurred (**Figure 2C**).

**Figure 2.**
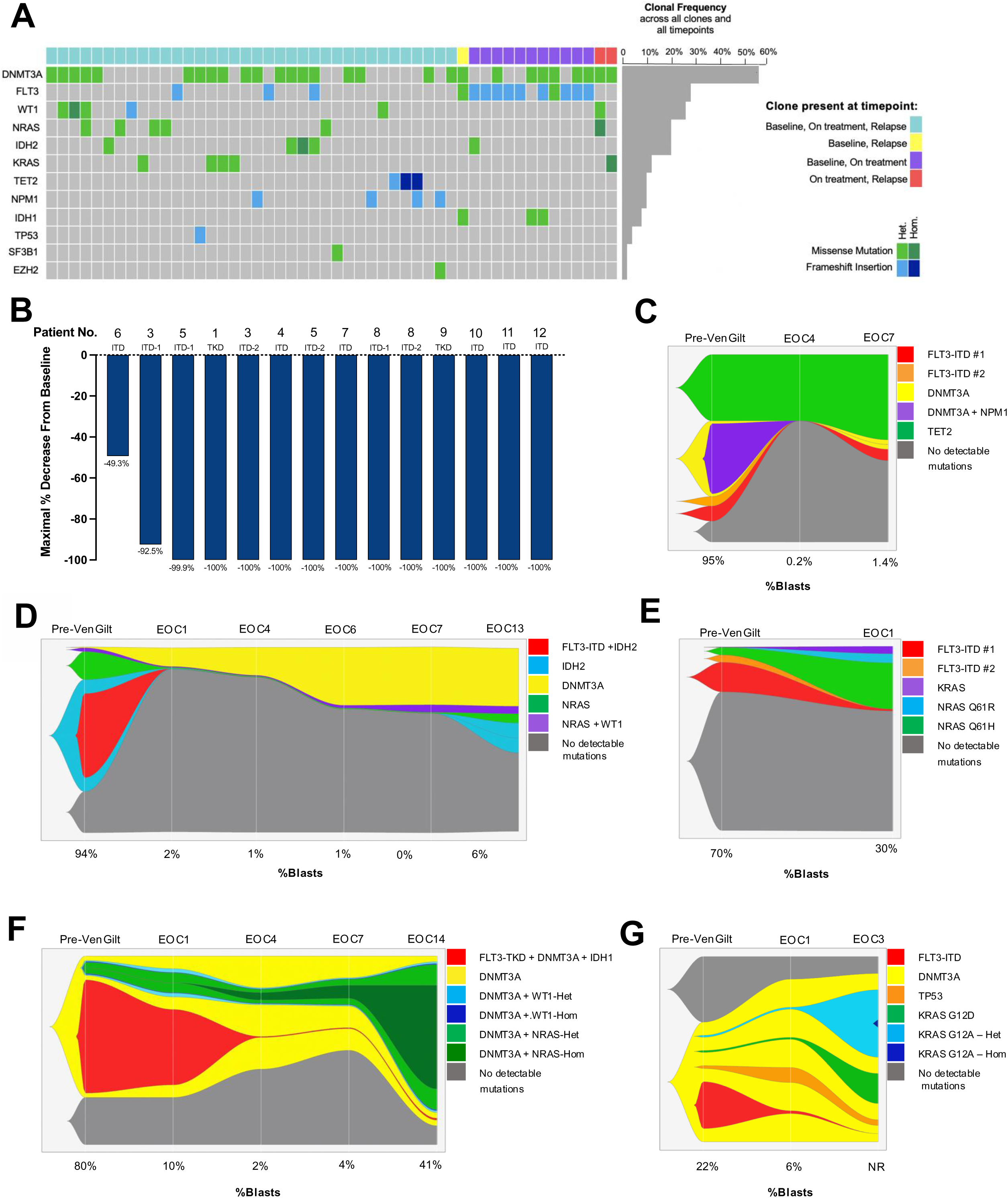
Diverse *FLT3*-mutated clones become undetectable with Ven/Gilt. **A.** Oncoprint of 50 genetically-defined clones across 12 patients and multiple timepoints. Each column is a unique clone, and clones are coded on the top row based on the clinical timepoint in which they were detected. Mutations are color-coded based on type of mutation and zygosity. **B.** Waterfall plot of 14 *FLT3-*mutated clones across 12 patients. Clones are coded based on patient number and type of *FLT3* mutation present (Internal Tandem Duplicate, ITD; Tyrosine Kinase Domain, TKD). *Select fishplots illustrate dynamic clonal architecture as measured by single cell (SC) analysis. For each plot, top x-axis indicates clinical timepoint (EOC, end of cycle) and bottom x-axis indicates percent blasts on corresponding clinical bone marrow biopsy, if available (NR, Not Reported)*. **C.** 3,655 SCs from Patient 5. In Patient 5, 1 *FLT3-*mut clone was eradicated while 1 initially decreased, then later recurred, on therapy. **D.** 30,550 SCs from Patient 2. In Patient 2, the *FLT3/IDH2* co-mutated clone was eradicated; *IDH2, NRAS,* and *WT1*-mutated clones later recurred. **E.** 5,606 SCs from Patient 3. In Patient 3, 1 *FLT3-*mut clone was eradicated while 1 decreased in size, but remained detectable. Diverse *N-* and *KRAS*-mutated clones expanded on therapy. **F.** 14,435 SCs from Patient 1. In Patient 1, the *FLT3*-mutated clone decreased in size on therapy while the *NRAS*-mutated clone was selected for. **G.** 9,571 SCs from Patient 4. In Patient 4, the *FLT3-*mutated clone was eradicated; multiple *KRAS*-mutated populations became dominant at relapse.

Four patients harbored clones in which FLT3 was co-mutated with IDH1/2, and in 3/4 patients, Ven/Gilt was active against the FLT3-IDH1/2 co-mutant clone (**Figure 2D, F; Figure S3B**). Thus, the presence of an IDH1/2 mutation in the same single cell as a FLT3 mutation did not obligate resistance to Ven/Gilt therapy, though IDH1/2 and FLT3 co-mutant clones have been implicated in resistance to both enasidenib and ivosidenib^11,23^. Interestingly, Patient 2 (**Figure 2D**) did show expansion of an IDH2-mutated clone at relapse, although this clone did not have a FLT3 co-mutation. Taken together, these findings suggest that in contrast to IDH1/2 inhibitors, FLT3 and IDH1/2 co-mutant clones may remain sensitive to Ven/Gilt. Ven/Gilt targeted therapy was also active against clones without *FLT3* mutations. 20 clones without *FLT3* mutations (in 11 individual patients), including clones with *IDH1, IDH2, DNMT3A, NPM1, TET2, EZH2, WT1,* and *NRAS* mutations, decreased by > 10% on treatment. This suggests that in addition to rapid activity against *FLT3*-mutated populations, Ven/Gilt was also active against some non-*FLT3-*mutated populations.

### Ven/Gilt selects for pre-existing RAS-mutated clones

Longitudinal SC DNAseq revealed that, of the 50 genetically distinct subclones detected across all timepoints, only 2 new clones appeared *de novo* on Ven/Gilt therapy **(Figure 2A**). All other clones were detected prior to Ven/Gilt therapy. This data suggests that, in this pre-treated population, genetic resistance was primarily driven by clonal selection rather than the development of new mutations. Interestingly, we did not observe development of or selection for any on-target resistance mutations within the *FLT3* gene within this clinical trial cohort, despite sequencing the entirety of *FLT3*.

Five patients had one or more pre-existing *N-* or *KRAS*-mutated clones (**Figure 2D-G; Figure S3A**). All heterogenous *RAS*-mutated populations were detectable prior to Ven/Gilt therapy as minor subclones, ranging from 0.001% - 18.1%. Notably, in 2 patients, these small, pre-existing *RAS-*mutated populations were not detectable by bulk sequencing (**Figure 1C**). In 4 patients, 9 pre-existing *N/KRAS*-mutated clones expanded on targeted therapy; all 4 of these patients had prior TKI therapy (**Figure 2D-G**). The kinetics of these *RAS*-mutant subclones was variable, driving disease relapse in 1-3 treatment cycles in some patients (**Figure 2E, G**), and up to 13-14 cycles in others (**Figure 2D, F**). Only 1 patient had a pre-existing *RAS*-mutated clone which did not expand on therapy; interestingly, that patient also had not received prior TKI therapy (**Figure S3A**). Out of all 50 clones detected across all patients and timepoints, only 2 new clones developed *de novo* on therapy. Notably, both new clones were homozygous for *RAS* mutations (**Figure 2A**) and developed in patients who harbored pre-treatment clones heterozygous for the same *RAS* mutations (**Figures 2F-G**), suggesting a strong selective pressure for increased RAS signaling in these patients.

### AML clones upregulate monocytic cell surface markers on Ven/Gilt

Using DAbseq, we next sought to measure the association between specific genotypes and immunophenotypes and determine how this association changed over time. Eight patients had longitudinal SC multi-omic profiling available (**Figure 3A**). With treatment, we noted shifts in genotype-immunophenotype association across the cohort of patients with available serial samples: *NRAS*-mutated cells became significantly associated with expression of monocytic markers CD11b and CD13 (Point-biserial correlation coefficient 0.76; p < 0.001 and 0.49; p = 0.005, respectively) and *IDH2*-mutated cells became significantly associated with CD11b (p = 0.035) **(Figure 3A**). **Figures 3B** and **S4** detail immunophenotypic evolution of 18 clones across 7 patients with longitudinal multi-omic profiling; only genotypes present in ≥ 2 clones are included. Across 7 *N/KRAS*-mutated clones, normalized median expression of monocytic markers CD11b and CD13 significantly increased with treatment (p = 0.047 and 0.006, respectively) with concurrent decrease in CD4 and CD45 (p = 0.034 and 0.023). This pattern was also observed in non-*RAS*-mutated genotypes. In 5 *DNMT3A*-mutated clones without co-mutations, expression of CD11b and CD13 significantly increased (p = 0.048 and p = 0.039, respectively); similar patterns were observed in 2 *IDH2*-mutated clones as well. By contrast, 3 *FLT3*-mutated clones did not demonstrate a significant change in CD11b or CD13.

**Figure 3.**
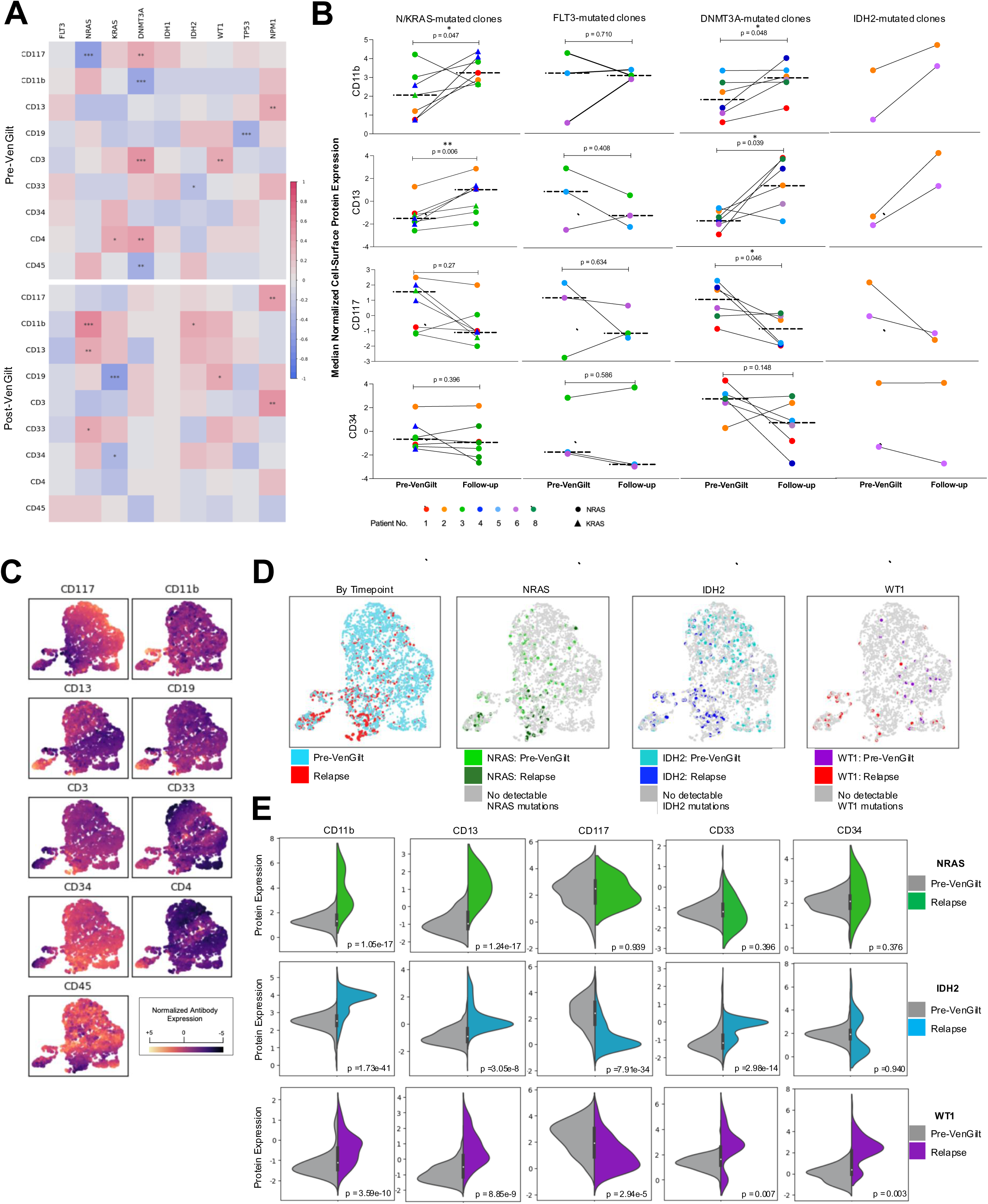
Multiomic SC sequencing reveals both genetic and immunophenotypic clonal evolution with Ven/Gilt. **A.** Correlation matrix of association between mutations and 9 cell-surface proteins in 6 patients with paired pre-Ven/Gilt (top) and post-Ven/Gilt (bottom) samples. Patients 1, 2, 3, 4, 5, 6, 8, and 10 are represented and the genes shown are reflective of those mutated in ≥ 2 clones. The plot shows co-occurrence (red) or exclusivity (blue) based on point-biserial estimate with statistician significance indicated as *p , .05; **p , .01; ***p , .001. **B.** Dot plot comparing pre-Ven/Gilt and post-Ven/Gilt samples from patients with *NRAS, KRAS, FLT3, DNMT3A,* or *IDH2*-mutated clones. The y-axis is median normalized expression of CD11b, CD13, CD117, and CD34 cell-surface proteins. Dashed lines indicate medians and dots are color and shape-coded based on source patient and/or mutation type. Significance from paired T tests indicated as *p, .05; **p , .01. **C.** Immunophenotype-derived UMAP from 6,512 integrated from pre-treatment and relapse timepoints from Patient 2. Cells are color-coded based on expression of cell-surface immunophenotypic proteins. **D.** UMAP from Panel C. Cells are color-coded based on time point, *NRAS* mutation plus time point, *IDH2* mutation plus timepoint, and *WT1* mutation plus timepoint. **E.** Split violin plot comparing expression of CD11b, CD13, CD117, CD33, and CD34 for cells pre-Ven/Gilt (gray) vs at relapse (green, blue, or purple) for *NRAS* (top row), *IDH2* (center row), and *WT1* (bottom row) mutated subclones.

Although RAS mutations have been associated with monocytic differentiation in prior studies^18^, we observed a heterogenous association between RAS mutant status and monocytic differentiation in pre-treatment clones (**Figure 3A, S2D-F, S4-5**). After treatment with Ven/Gilt, however, there was a strong association between RAS-mutant genotypes and upregulation of CD11b and CD13, which was particularly striking in select patients (**Figure 3A, C-E, Figure S5-6**). Interestingly, in Patient 1, who harbored both heterozygous and homozygous *NRAS*-mutant populations, the degree of increased CD11b/13 expression was greater in the *NRAS-*homozygous subclone (**Figure S6 C-E**). This could suggest a potential dose-dependent effect of oncogenic RAS signaling on expression of monocytic markers. Although this pattern of CD11b and CD13 upregulation was most evident in RAS mutant clones, we also observed upregulation of monocytic markers in diverse non-RAS mutant clones as well, including clones harboring *IDH2, WT1*, and *DNMT3A* mutations (**Figure 3C-E; Figure S6**), suggesting that monocytic selection and/or evolution can occur across genotypes.

### Individual patients contain transcriptionally distinct immature and monocytic leukemic populations which are dynamic on Ven/Gilt therapy

To further investigate the association between AML differentiation state, RAS signaling, and therapeutic resistance, we performed SC Cellular Indexing of Transcriptomes and Epitopes by Sequencing (CITE)seq on available paired pre-Ven/Gilt and relapse samples from four patients with RAS-mutated clones that expanded on therapy, for a total of 42,853 single cells (median 5,357 cells per time point). This included 2 patients treated on trial (Patients 1, 2) and 2 patients treated off-trial with identical dosing of both agents, all who had evidence of RAS-mutated clonal expansion based on bulk DNA sequencing (Patients 13, 14; **Table S1**). Of note, while the 2 off-trial patients had clonal expansion of RAS mutations with relapse, unlike the clinical trial cohort, they also each had expansion of an on-target mutation within the *FLT3* gene (*FLT3* N701K and F691L), a pattern not observed in any on-trial patient (**Figure S7**).

We employed multiple orthogonal methods to annotate and classify leukemia cell phenotype using transcriptomic and immunophenotypic data. We applied the RNAseq-based classification scheme clustifyr, as per previous work annotating leukemic blasts from SC RNAseq data^24^ (**Figure S8A**) and examined co-expression of cell-surface markers which can distinguish blasts from normal hematopoietic stem and progenitor cells^25–28^ (**Figure S8B**). Single cells classified as “unassigned”, “CD34+”, “CD14+”, or “CD16+” by clustifyr *and* expressing at least 2 of CD33, CD123, HLA-DR, or CLEC12A were considered leukemic blasts. To subclassify leukemic blasts, we next calculated the cell state score of the six AML differentiation states first described by van Galen et al by analyzing SC transcriptome data of multiple AML samples; importantly, this classification schema compared AML blasts against normal hematopoietic cells, thus affirming our classification of leukemic blasts vs normal cells^29^ (**Figure S8C, Figure 4A**). From the resultant bimodal distribution, leukemic cells in the top 50th percentile for HSC- or Progenitor-like AML were classified as immature leukemia, while patients in the top 50th percentile for Monocyte-like AML were classified as monocytic leukemia (**Figure 4A**).

**Figure 4.**
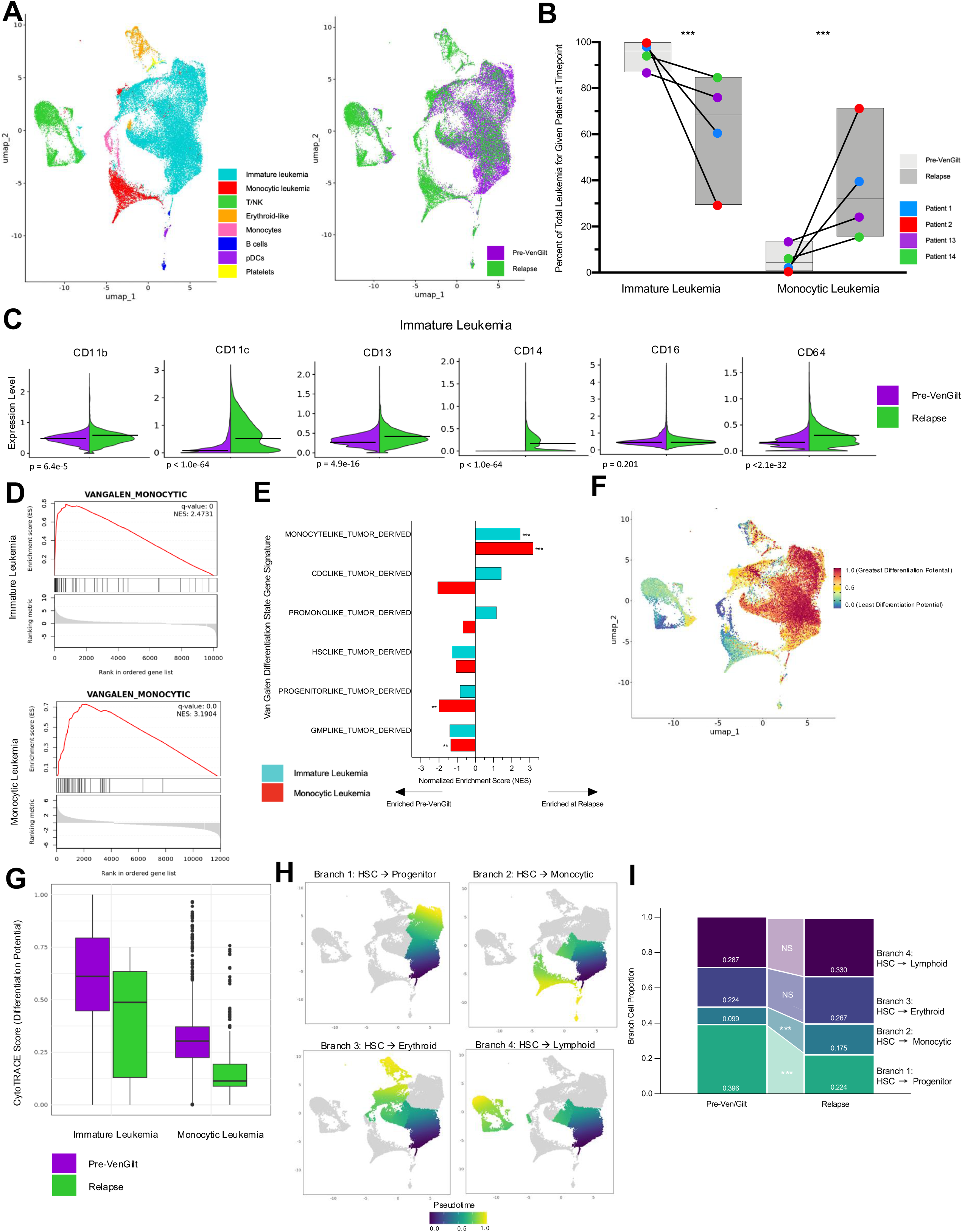
Immature and Monocytic Leukemia populations are dynamic under therapeutic pressure, and both become increasingly monocytic. **A.** RNA-derived UMAP from 42,853 single-cells from paired pre-treatment and relapse timepoints from 4 patients treated with Ven/Gilt. Left: Cells are color-coded based on cell type as identified via combination of RNA-based classifiers and cell-surface markers. Right: Cells are color-coded based on time point. **B.** Dot plot of monocytic and immature leukemia cells as a proportion of all leukemia cells for each time point. For each patient, the relative proportion of monocytic leukemia significantly decreased from pre-treatment to relapse and the relative proportion of immature leukemia increased, indicated as *** p < 0.001. **C.** Split violin plot comparing normalized cell surface antibody expression of monocytic markers between Pre-Ven/Gilt vs Relapse timepoint for the monocytic leukemia population. Horizontal lines indicate median expression. **D.** Enrichment profile and ranking metric score for the monocyte-like AML gene set in the immature (*top*) and monocytic (*bottom*) leukemia populations. For both populations, the gene set is significantly positively enriched at relapse relative to pre-Ven/Gilt. **E.** Bar plot of normalized enrichment scores (NES) from Gene Set Expression Analysis (GSEA) of immature (turquoise) and monocytic (red) leukemia cells. Positive scores indicate enrichment at relapse and negative scores indicates enrichment pre-Ven/Gilt. Statistical significance is indicated as ***q <0.001, ** q <0.01, *q <0.05. **F.** UMAP from 4A. Cells are color-coded based on cytoTRACE score from 0 (most differentiated) to 1 (least differentiated). **G.** Box and whiskers plot of cytoTRACE score for immature (*left*) and monocytic *(right*) leukemia populations pre-Ven/Gilt (purple) and at relapse (green). **H.** UMAP from 4A. Cells are color-coded based on psuedotime as determined by the Lamian computational framework. The 4 subpanels highlight cells comprising the 4 primary trajectories (1) HSC → Progenitor, (2) HSC → Monocytic, (3) HSC → Erythroid, and (4) HSC → Monocytic. **I.** Bar plot depicting changes in branch cell proportion between pre-Ven/Gilt vs. Relapse. From pre-Ven/Gilt to relapse, the HSC → Progenitor branch significantly decreased, the HSC → Monocytic branch significantly increased, and the HSC → Lymphoid and HSC → Erythroid branches did not significantly change. Statistical significance is indicated as *** p <0.001; NS: non-significant.

We used unbiased single-cell gene set enrichment analysis (GSEA) using the molecular signature database (mSigDB) hallmark and C2 gene sets^30,31^ as well as gene sets associated with specific AML differentiation states^29,32^, to determine differences in transcriptional profiles between these two AML subpopulations. At baseline prior to Ven/Gilt therapy, the common monocytic and immature leukemia clusters demonstrated distinct transcriptional profiles (**Figure S9A-B; Table S5**). Relative to the monocytic leukemia population, the immature leukemia population demonstrated significant enrichment for stem-cell associated gene sets and gene sets associated with HSC and progenitor-like AML. By contrast, monocytic leukemia cells were enriched for monocyte-like AML as well as an MLL (KMT2A)-rearranged leukemia signature, previously shown to be enriched in monocytic AML^24^, further affirming our leukemia classification schema (**Figure S9B**). These findings are consistent with those of Pei et al^17^ who noted similar distinct transcriptional profiles associated with monocytic vs primitive AML leukemic stem cells (LSCs). Notably, consistent with prior observations linking RAS activation and monocytic differentiation, the monocytic leukemia population was also enriched for a RAS activation signature (**Figure S9B**).

Of the 4 patients analyzed, all contributed to both the monocytic leukemia and immature leukemia clusters (**Figure S8D**). All patients demonstrated significant shifts in the relative proportion of the monocytic and immature leukemia populations over time. On an individual patient level, for each of the 4 patients the proportion of immature leukemia cells decreased on therapy while the proportion of monocytic leukemia cells increased (median percent change 21.4%, range 9.4% - 70.5%) (**Figure 4B**). Taken together, this analysis demonstrates that in individual patients, pre-existing monocytic leukemia populations were selected by Ven/Gilt treatment.

### Immature and monocytic AML clones increase differentiation and monocytic features on Ven/Gilt

In addition to changing in relative proportion, the immature and monocytic leukemia populations also demonstrated dynamic changes in both immunophenotype and gene expression profile on Ven/Gilt therapy. As expected, at baseline, the immature leukemia and monocytic leukemia populations demonstrated distinct immunophenotypes, with the monocytic leukemia population having significantly higher expression of multiple monocytic markers (**Figure S8E**).

With Ven/Gilt treatment, monocytic population further increased expression of 3 monocytic markers CD11b (0.77 vs 0.92, p <3.3e-47), CD13 (0.34 vs 0.42, p <1.9e-21), and CD14 (0.29 vs 0.52, p=1.7e-11) while CD11c, CD16, and CD64 remained unchanged (**Figure S8F**). Surprisingly, the immature leukemia population significantly increased expression of all assessed monocytic markers with the exception of CD16, including CD11b (pretreatment vs relapse: 0.47 vs 0.51 normalized expression units, p = 6.4e-5), CD11c (0.13 vs 0.57, p <1.0e-50), CD13 (0.31 vs 0.38, p <4.9e-16), CD14 (0.02 vs 0.18, p<1.0e-50) and CD64 (0.16 vs 0.23, p <2.1e-32) (**Figure 4C**).

As an alternative metric for assessing differentiation state, we applied CytoTRACE [for cellular (Cyto) Trajectory Reconstruction Analysis using gene Counts and Expression] to our single cell transcriptional dataset. CytoTRACE is a computational framework for predicting the differentiation potential of a single cell based on transcriptional data about numbers of expressed genes, covariant gene expression, and local neighborhoods of transcriptionally similar cells. CytoTRACE provides a score for each cell representing its stemness within a given dataset, ranging from 0 to 1, with higher scores indicating greater stemness^33^. When applied to our cohort, we found that the pre-treatment differentiation potential of the monocytic population was lower than that of the immature population, as expected given our cell classification framework (**Figure 4F**). With Ven/Gilt treatment, the differentiation potential of both populations significantly decreased, indicating evolution to a more differentiated and less stem-like cell state (Immature leukemia: median CytoTRACE 0.60 pre-Ven/Gilt vs 0.54 at relapse, p < 2.0e-50; Monocytic leukemia: median CytoTRACE 0.33 pre-Ven/Gilt vs 0.17 at relapse, p < 2.0e-50 (**Figure 4G**), consistent with our observation that both cell types adopt a more differentiated monocytic cell state under treatment selection.

### Pseudotime analysis identifies enrichment for a monocytic differentiation trajectory at relapse relative to pre-treatment

To further interrogate how differentiation trajectories change from pre-treatment to relapse, we performed pseudotime analysis on all integrated cells using Lamian, a computational framework for pseudotime analysis using SC data from both multiple biological samples and multiple conditions or timepoints^34^. Using Lamian, we identified 4 primary trajectories: (1) HSC → Progenitor, (2) HSC → Monocytic, (3) HSC → Erythroid, and (4) HSC → Lymphoid (**Figure 4H**).

We conducted repeated bootstrap sampling of cells along these trajectories to calculate branch proportions at pre-Ven/Gilt vs relapse timepoints. Branch proportions differed significantly for the 2 trajectories predominantly occurring within the leukemic populations, HSC → Progenitor (0.396 vs 0.224 for pre-Ven/Gilt vs Relapse, p < 0.001) and HSC → Monocytic (0.099 vs 0.175 for pre-Ven/Gilt vs Relapse, p < 0.001), again indicating increased enrichment for monocytic differentiation at relapse. By contrast, the HSC → Lymphoid and HSC → Erythroid trajectories, which terminated in non-leukemic populations, did not significantly differ between timepoints (HSC → Lymphoid: 0.224 vs 0.267, p = 0.23; HSC → Erythroid: 0.297 vs 0.320, p = 0.73 for pre-Ven/Gilt vs relapse, respectively) (**Figure 4I**).

### Monocytic gene expression and venetoclax resistance signatures are enriched in both immature and monocytic AML subpopulations at relapse

We next performed GSEA to compare both leukemic subpopulations to identify transcriptional programs associated with monocytic differentiation which could be underlying Ven/Gilt resistance. Prior work identified changes in a gene signature associated with venetoclax resistance in AML which includes genes associated with anti-apoptotic proteins BCL2, MCL1, and BCL2A1^16^. Expression of these genes differed between leukemia populations at baseline. At baseline prior to Ven/Gilt therapy, expression of BCL2 was significantly higher in the immature leukemia population, while expression of MCL1, BCL2A1, BID, BAK1, B2M, and BAK1 were significantly higher in the monocytic population, consistent with prior work^17^ (**Figure S10A**). When assessed via GSEA, pre-treatment, the venetoclax resistance signature was also significantly enriched in the monocytic population relative to the immature population (NES 1.77, p = 0.007) (**Figure S9B**), supporting prior observations that monocytic AML subclones demonstrate intrinsic resistance to venetoclax^17^.

Using GSEA, we noted dynamic shifts in transcriptional profiles of both the immature and monocytic leukemia populations with treatment (**Figure S9C-D; Table S6-7**). Mirroring the cell-surface protein data, both populations demonstrated significant enrichment for a monocyte-like AML transcriptional signature with resistance to targeted therapy (Immature cluster: normalized enrichment score [NES] 2.47, q-value 0.0; Monocytic cluster: NES 3.19, false discovery rate [FDR] q-value 0.0;) (**Figure 4D; Figure S9E**). No other gene sets derived from a specific AML differentiation state were significantly positively enriched in either leukemia type, although progenitor-like and GMP-like AML signatures were negatively enriched in the monocytic leukemia population after treatment (**Figure 4E**).

With Ven/Gilt treatment, genes associated with venetoclax resistance demonstrated distinct evolution in the immature vs monocytic leukemia populations (**Figure S10B-C**) with Ven/Gilt resistance. The immature leukemia populations demonstrated a significant decrease in BCL2 (p = 2.4e-4), MCL1 (p = 2.0e-68) and CD34 (p<1.0e-99) expression, while expression of BID (p = 3.0e-31) and CD68 (p < 1.0e-99) significantly increased. By contrast, the monocytic leukemia population demonstrated a significant decrease in FUT4 (p = 3.6e-13) and increase in MCL1(p = 4.6e-44), ITGAM/CD11b (p = 1.3e-29), CD14 (p = 9.6e-12), and CD36 (9.3e-4). Both populations showed marked increase in expression of BCL2A1, an anti-apoptotic previously associated with venetoclax resistance^16^ (immature: p = 3.2e-24; monocytic: p = 4.5e-29) with resistance (**Figure 5A, Figure S10B)**. When assessed via GSEA, the venetoclax resistance signature become modestly enriched in the immature population (NES for relapse vs pre-treatment 1.45, p = 0.04), while expression of this signature in the monocytic population remain unchanged (NES 0.62, p = 0.90) (**Figure S9C-D**). Overall, these changes suggest an overall shift in anti-apoptotic gene expression for both leukemia populations, consistent with increased venetoclax resistance.

**Figure 5.**
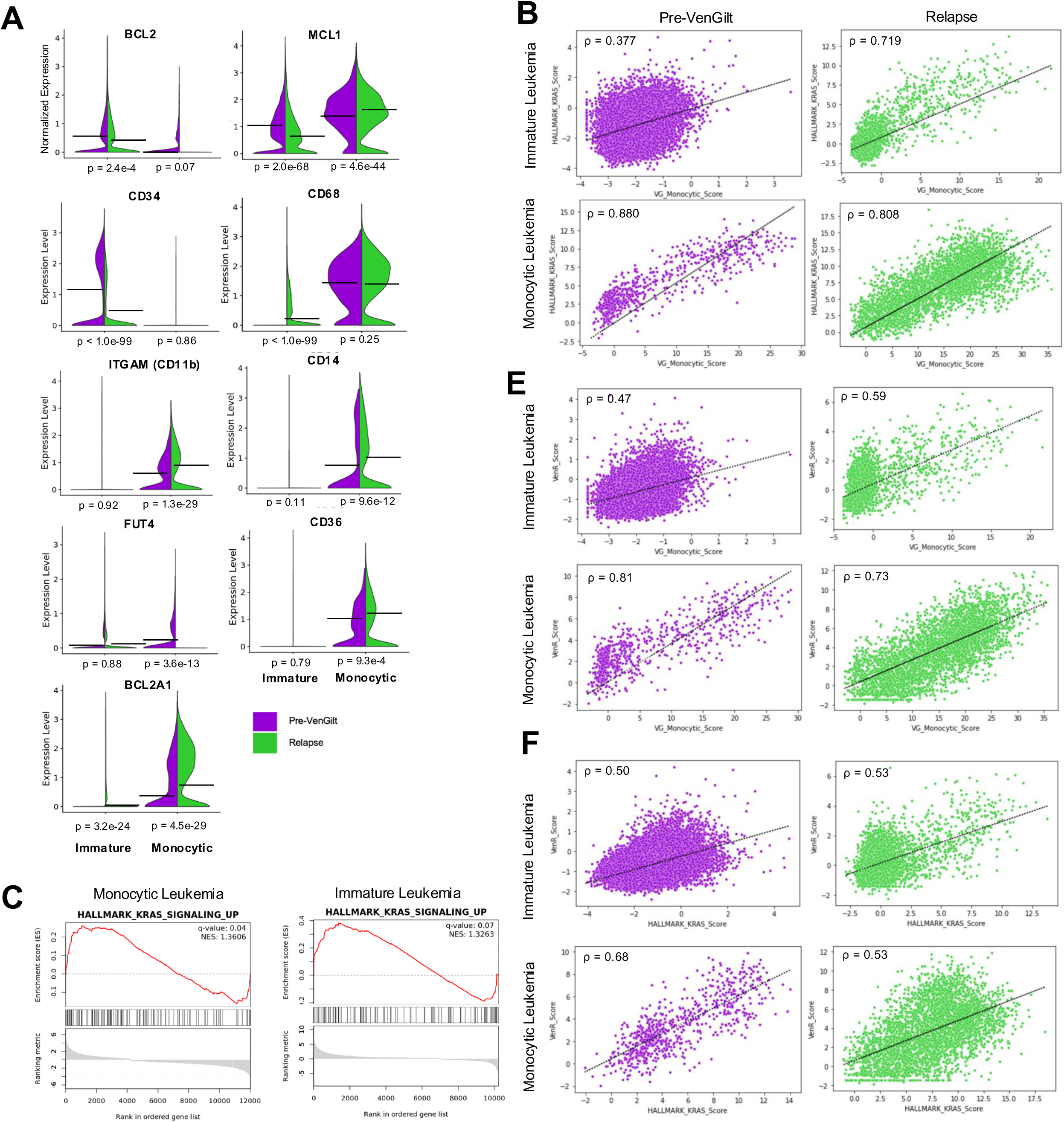
RAS signaling increases with therapeutic pressure; transcription signatures of both RAS signaling and venetoclax resistance genes are associated with monocytic cell state. **A.** Split violin plot comparing expression of select genes associated with venetoclax resistance in the literature in the immature vs monocytic leukemia populations at baseline and prior to Ven/Gilt therapy. **B.** Scatterplot depicting correlation between RAS signaling vs monocytic cell state in the immature and monocytic populations at baseline and relapse. Each point represents a single cell. **C.** Enrichment profile and ranking metric score for RAS signaling in the monocytic (*left)* and immature (*right*) populations. In the monocytic population, RAS signaling is significantly positively enriched at relapse relative to pre-Ven/Gilt. **E.** Scatterplot depicting correlation between venetoclax resistance vs monocytic cell state in the immature and monocytic populations at baseline and relapse. Each point represents a single cell. **F.** Scatterplot depicting correlation between venetoclax resistance vs RAS signaling in the immature and monocytic populations at baseline and relapse. Each point represents a single cell.

Prior studies have also implicated specific gene signatures, including AML with MLL fusions, oxidative phosphorylation, and purine metabolism, in association with monocytic AML relapsed or refractory to venetoclax-containing therapies such as venetoclax/azacitidine^17,24^. In our data, a MLL fusion gene signature was enriched in monocytic leukemia and with resistance in the immature, but not the monocytic, cell population (Immature population: NES 1.71, p = 0; Monocytic population: NES 0.88, p = 0.71). However, signatures associated with purine metabolism and oxidative phosphorylation were not enriched in monocytic compared to immature leukemia prior to treatment or at relapse in either leukemia subtype (**Figure S9D-E**). These findings are distinct from these earlier findings which associate these signatures with monocytic AML and venetoclax resistance^17,24^. Cumulatively, these findings suggest that while mechanisms of resistance to venetoclax may demonstrate some overlap, resistance signatures may be unique to particular venetoclax combinations.

### RAS-associated transcriptional activation increases with Ven/Gilt treatment and is tightly associated with a monocytic cell state

Given the known association of RAS signaling with monocytic differentiation^19^, we investigated how RAS-driven transcriptional activation correlates with AML differentiation state and changes with Ven/Gilt treatment. Across all four patients, the integrated monocytic leukemia population demonstrated a significant enrichment for genes associated with RAS signaling activation with Ven/Gilt treatment (NES 1.36, q = 0.04) (**Figure 5C**). We measured the association between RAS signaling and monocytic cell state at a transcriptional level by calculating the expression score of both the Hallmark KRAS signaling upregulation^31^ and monocyte-like AML^29^ gene sets, using a methodology similar to that used to apply gene set expression scores to bulk RNAseq data^18,35^. Of note, only 2 of > 300 genes are shared between the two analyzed gene sets.

At baseline prior to Ven/Gilt, the immature leukemia population demonstrated low RAS score, low monocyte-like AML scores (median -1.27 and -2.04, respectively) and a weak correlation between RAS signaling and monocyte-like AML (Spearman ρ = 0.377) (**Figure 5B**). By contrast, pre-Ven/Gilt, the monocytic leukemia population demonstrated higher scores for both RAS signaling and monocyte-like AML (median 4.31 and 3.30, respectively) and a markedly stronger correlation (Spearman ρ = 0.880), which persisted at time of relapse (Spearman ρ = 0.808). Interestingly, though none of these patients acquired new RAS mutations at relapse, both RAS signaling and monocyte-like AML scores increased at relapse in the immature leukemia population (median -0.43 and -1.77, respectively), and this increase was tightly correlated (Spearman ρ = 0.791). Taken together, these findings demonstrate that activated RAS signaling and monocytic differentiation are enriched and tightly correlated in monocytic leukemia cells as well as in immature leukemia cells at relapse on Ven/Gilt treatment. We also investigated the relationship between known venetoclax resistance genes, monocytic cell state, and transcriptional signatures of activated RAS signaling. The relationship between venetoclax resistance and monocytic differentiation score was correlated in both immature and monocytic leukemia, most pronounced in the monocytic leukemia population, and this was unchanged at relapse (**Figure 5E**) (Spearman correlation coefficient 0.81 pre-Ven/Gilt, 0.73 at relapse). The venetoclax resistance score was similarly correlated with the RAS signaling score as well, albeit to a lesser extent (**Figure 5F**)

### Distinct molecular mechanisms are associated with resistance to venetoclax and venetoclax combinations

While multiple mechanisms of venetoclax resistance have been described, the mechanistic underpinnings of resistance to individual venetoclax containing-combinations remain relatively unknown. While data derived from Ven/Gilt-treated patients suggest both RAS signaling and monocytic differentiation are associated with resistance to Ven/Gilt, we found that gene signatures previously associated with venetoclax resistance, such as oxidative phosphorylation^17^, were not uniformly associated with Ven/Gilt resistance. To better understand distinct molecular mechanisms associated with venetoclax vs venetoclax/gilteritinib resistance, we cultured *FLT3-ITD* and *NRAS* co-mutant Molm14 cell lines in escalating doses of drug to develop cell lines resistant to venetoclax (Ven-R) and venetoclax/gilteritinib (Ven/Gilt-R) combination (**Figure 6 A-B)**.

**Figure 6.**
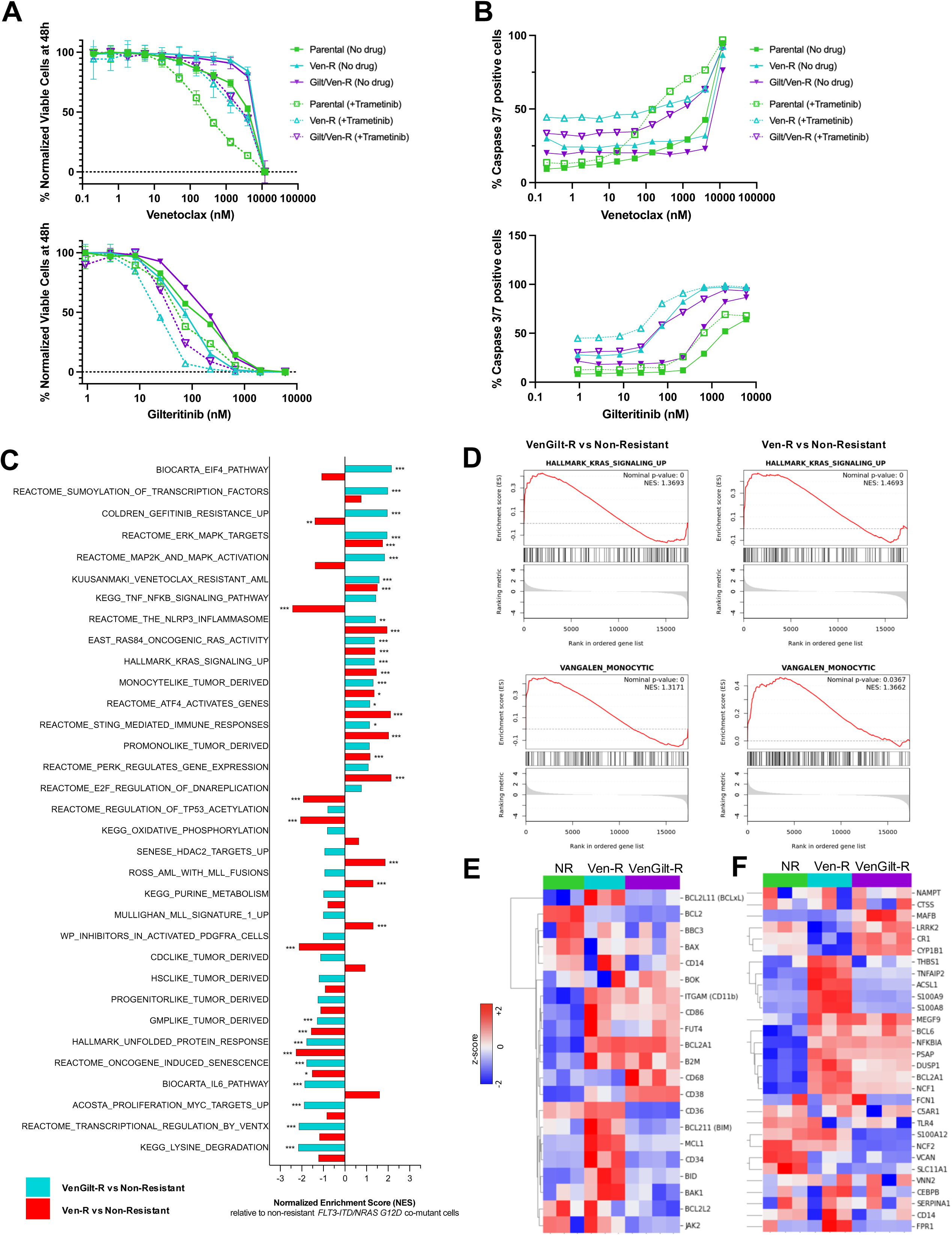
Targeted therapy resistance is associated with monocytic differentiation and increased RAS signaling *in vitro*. **B.** Heatmap of normalized gene expression of select genes associated with Venetoclax resistance in the non-resistant (NR), Ven-R, and Ven/Gilt-R MOLM-14 (*FLT3-*ITD) and *NRAS* G12C co-mutant cell lines. **C.** Heatmap of normalized gene expression of select genes associated with monocytic differentiation in the non-resistant (NR), Ven-R, and Ven/Gilt-R MOLM-14 (*FLT3-*ITD) and *NRAS* G12C co-mutant cell lines. **D.** Dose response curves representing relative proliferation of Ven-R, Ven/Gilt-R, and parental non-resistant *NRAS* G12C mutant MOLM-14 (*FLT3*-ITD+) cells with and without the addition of trametinib after 48 hours of increasing doses of venetoclax (top) and gilteritinib (bottom). Experiments done in 3 technical replicates and error bars represent standard deviation **E.** Dose response curves representing caspase 3/7 expression of Ven-R, Ven/Gilt-R, and parental non-resistant *NRAS* G12C mutant MOLM-14 (*FLT3*-ITD+) cells with and without the addition of trametinib after 24 hours of increasing doses of venetoclax (top) and gilteritinib (bottom). Experiments done in 3 technical replicates and error bars represent standard deviation. **F.** Bar plot of normalized enrichment scores (NES) from Gene Set Expression Analysis (GSEA) comparing venetoclax/gilteritinib-resistant (Ven/Gilt-R) vs non-resistant *FLT3-ITD/NRAS* co-mutant cells (turquoise) and venetoclax-resistant (Ven-R) vs non-resistant *FLT3-ITD/NRAS* co-mutant cells (red). Positive scores indicate enrichment in the resistant cells. Statistical significance is indicated as ***q <0.001, ** q <0.01, *q <0.05. **G.** Enrichment profile and ranking metric score for the RAS signaling (*top)* and monocyte-like AML (*bottom*) gene sets in the Ven/Gilt-R (*left)* and Ven-R *(right)* cells vs the non-resistant cells.

Bulk RNAseq analysis of the 3 cell lines (*FLT3-ITD/NRAS* non-resistant, *FLT3-ITD/NRAS* Ven-R, and *FLT3-ITD/NRAS* Ven/Gilt-R) demonstrated distinct transcriptional signatures (**Figure S11**). By GSEA, relative to non-resistant cells, Ven/Gilt-R cells demonstrated significant enrichment for gene signatures associated with venetoclax resistance (NES = 1.59, p = 0.0), RAS signaling (NES = 1.37, p = 0.0), and monocytic differentiation (NES = 1.32, p = 0.0), mirroring what we observed with longitudinal patient samples **(Figure 6C-D; Table S8-9)**. Similarly, Ven-R cells also demonstrated significant enrichment for venetoclax resistance (NES = 1.51, p = 0.0), RAS signaling (NES = 1.47, p = 0.0), and monocytic differentiation (NES = 1.37, p = 0.0) signatures.

Both Ven/Gilt-R and Ven-R resistant cell lines demonstrated transcriptional upregulation of the monocytic marker ITGAM/CD11b, again mirroring both our patient DAbseq and CITEseq data **(Figure 6E)**. In parallel, the cell lines showed an increase in cell surface protein expression of CD11b (**Figure S12A**). Transcriptional upregulation of CD86, BCL2A1, and B2M and downregulation of BCL2 was also observed in both resistant cell lines. However, many venetoclax-resistance associated genes, including BID, BAK2, BCL2L11, BCL2L1, and MCL1, had greater transcriptional upregulation in Ven-R compared to Ven/Gilt-R cells, potentially suggesting distinct resistance mechanisms. Similarly, while both Ven-R and Ven/Gilt-R cells demonstrated increased monocytic and RAS transcriptional signatures, the individual gene expression patterns differed between the two cell lines (**Figure 6E-F, Figure S11A**). Western Blot analysis confirmed increased ERK and AKT phosphorylation in both resistant cell lines, consistent with increased RAS signaling activation (**Figure S12B**).

While RAS signaling activation and a RAS activation signature were upregulated in both resistant cell lines, bulk whole exome sequencing uncovered distinct mutational profiles. In Gilt/Ven-R cells, we identified point mutations in the RAS effector RAF1 S257L (48% VAF) and apoptotic effector BAX R94* (16% VAF). RAF1 S257L is a known activating mutation, associated with Noonan’s syndrome^36^ (nd likely accounts for upregulation of RAS signaling programs in this cell line. Additionally, BAX R94* is a known venetoclax-resistance mutation in AML^37^. However, in Ven-R cells, neither mutation was identified. Beyond these mutations, no additional mutations or copy number changes in the RAS/MAPK signaling pathway, genes implicated in venetoclax resistance, or other significant oncogenic drivers were identified in either model (**Table S10-11**). Taken together, this analysis further suggests that both RAS activation and monocytic differentiation are enriched with resistance to venetoclax and venetoclax/gilteritinib. These data further suggest that increased activation of RAS signaling and monocytic differentiation can occur via both genetic and non-genetic mechanisms. Beyond RAS signaling activation, gene expression programs previously associated with venetoclax resistance driven by monocytic AML differentiation, including oxidative phosphorylation and purine metabolism^17,24^ were not upregulated in either cell line.

### BCL2A1 overexpression drives resistance to venetoclax but not to gilteritinib

Given that BCL2A1 expression was uniformly upregulated in Ven/Gilt-treated patients and in both VenR and Ven/GiltR cell lines, and that BCL2A1 overexpression has previously been identified to confer resistance to venetoclax^38^, we sought to determine the impact of BCL2A1 overexpression on response to gilteritinib and Ven/Gilt. Using a doxycycline-inducible system, we generated *BCL2A1* overexpressing MOLM-14 cells (**Figure S12C**). Induction of *BCL2A1* overexpression increased resistance to gilteritinib and Ven/Gilt (**Figure S12D**). These data confirm that BCL2A1 overexpression is a mechanism of resistance to Ven/Gilt and suggests that BCL2A1 may be a potential therapeutic target to overcome resistance to select venetoclax combinations such as Ven/Gilt.

### MEK inhibition resensitizes resistant cells to both to venetoclax and venetoclax combinations

Given evidence of increased RAS signaling and RAS transcriptional activation in both Ven-R and Ven/Gilt-R cells, we hypothesized that inhibition of downstream RAS signaling would resensitize resistant cells to both venetoclax and gilteritinib. In confirmation of this hypothesis, treatment with the MEK1/2 inhibitor trametinib restored sensitivity of both Ven/Gilt-R and Ven-R resistant cell lines to venetoclax and gilteritinib as measured by both proliferation and apoptosis (**Figure 6A-B**). In the case of gilteritinib, trametinib treatment actually sensitized both Ven/Gilt-R and Ven-R cell lines beyond the level of the parental line. Downregulation of ERK phosphorylation was associated with decrease in MCL1 and BCL2A1 levels in both resistant cell lines (**Figure S12B**), consistent with resensitization to venetoclax through MAPK pathway inhibition. Taken together, these results indicate that resistance to venetoclax and gilteritinib in both Ven/Gilt-R and Ven-R cells depends on RAS signaling activation. Moreover, these data suggest that RAS pathway inhibition effectively overcomes resistance to Ven/Gilt associated with monocytic differentiation and transcriptional RAS pathway activation, even in the absence of acquired RAS pathway mutation in Ven-R cells.

### Dynamic cell state and monocytic differentiation are observed with resistance in an independent and heterogeneous patient cohort

A strength of our single-cell analyses is the uniform nature of our patient cohort, all of which were treated with Ven/Gilt therapy and nearly all on the same phase 1b clinical trial. However, the extent to which similar changes in AML differentiation state occur in AML patients treated with other therapies is unknown. To investigate this, we analyzed bulk RNAseq data from 44 adult AML patients with paired pre- and post-treatment data available from the BEAT AML initiative^39^. This cohort is diverse, with 14 patients receiving cytotoxic chemotherapy and 17 various targeted kinase inhibitors, many on diverse clinical trials (**Table S12**); notably, no patients received gilteritinib and only 3 received venetoclax or venetoclax combinations.

Despite this heterogeneity, many patients demonstrated dynamic cell state with treatment, as measured by Van Galen differentiation state gene set scores (**Figure S13A**). Of the 17 patients treated with targeted therapies, there was a significant decrease in progenitor-like cell state (p = 0.039) and a trend toward increase in monocytic cell state (p = 0.15) with treatment (**Figure S13B**). In the cohort as a whole, monocytic differentiation correlated with RAS signaling activation (**Figure S13C**). In both the entire cohort and the subset of patients treated with targeted therapies, an increase in monocytic cell state with treatment tightly correlated with an increase in RAS signaling (all patients: *R =* 0.57; p = 5.0e-5; patients treated with targeted therapy: *R =* 0.74; p = 0.0007) (**Figure S13D**). While this cohort is highly heterogeneous, this analysis further confirms that monocytic cell state and RAS signaling are dynamic and tightly correlated, regardless of therapy.

## Discussion

AML is a heterogeneous disease characterized by genetic and immunophenotypic inter- and intra-tumor diversity. Across AML subtypes, BCL2 inhibition has emerged as an important component of AML therapy, and venetoclax combinations are now standard of care in both frontline and R/R setting across various genotypes^2,3^. For R/R FLT3-mutant AML specifically, Ven/Gilt combination therapy has potential to become standard of care. However, all patients on this combination eventually relapse absent stem cell transplantation, and understanding resistance mechanisms is critical to improve patient outcomes to this and other targeted therapies. In this context, we offer comprehensive correlative multiomic analyses of prospectively collected serial samples from all patients treated on the Ven/Gilt phase 1b trial with available samples^9^. To our knowledge, this is the first systematic multiomic single cell analysis from a uniform clinical trial cohort.

Our analysis reveals that response to Ven/Gilt occurs on the clonal, rather than the patient level. Clones within the same leukemia exhibit differential responses, highlighting that clonal selection drives both response and resistance. We find that Ven/Gilt effectively eradicates FLT3-mutant clones, with rapid decrease often observed over as little as 1 month of treatment. Notably, Ven/Gilt treatment also decreased clones *without* FLT3-mutation. This may be due to the effect of venetoclax alone, although it is also possible that elimination of crosstalk from FLT3-mutant clones results in death of non-FLT3 mutant clones. Future mechanistic work is needed to validate these theories. Importantly, SC DNA analysis reveals that the combination is effective even in FLT3 mutant clones with co-mutations (DNMT3A, IDH1, IDH2), and that co-mutational status can drive clonal resistance versus response. SC profiling of co-mutational status therefore has potential to direct therapeutic decision making in the era of multiple targeted therapies, such as in the case of FLT3 and IDH inhibitors^40^.

RAS mutations are an important cause of resistance to both FLT3^10,22^ and BCL2 inhibitors in AML^13^. Consistent with these findings, we demonstrate strong selection for RAS signaling activation in AML patients treated with Ven/Gilt combination treatment, including selection for pre-existing RAS mutant clones and in 2 cases, selection for new homozygous RAS mutant populations, suggesting that an increase in RAS signaling, beyond the presence of the mutation itself, drives resistance^41^. Despite the association of pre-existing RAS mutation with eventual relapse, the kinetics of relapse are challenging to predict, ranging from 1-13 cycles in our cohort. Except for 1 patient treated off-trial, in our cohort, RAS mutations were not co-mutated with FLT3 in resistant clones. This is potentially because FLT3-mutant clones are eradicated quickly with Ven/Gilt, prior to acquiring new resistance mutations. Perhaps for a similar reason, on-target FLT3 resistance mutations were rare, identified in only 2 patients treated off-trial, both of whom had prior exposure to gilteritinib (in contrast to patients treated on-trial, who were gilteritinib-naive). Of note, all patients in this cohort, both on- and off-trial, were heavily pre-treated, and the majority had received prior FLT3 TKIs. This may have selected minor, pre-existing RAS-mutant populations. Interestingly, the only patient with a RAS mutation that did not expand on Ven/Gilt had no history of prior TKI treatment, raising a question of whether prior TKI treatment may select for more resistant RAS mutant clones.

Though we can associate RAS mutations with Ven/Gilt resistance, RAS mutations alone are an imperfect clinical biomarker. At the time of relapse, we also show that RAS mutant (and some wild type) clones increase expression of monocytic cell surface markers. Using CITE-seq we show that all patients harbored both immature and monocytic AML clones prior to Ven/Gilt therapy. With treatment, pre-existing monocytic AML blast populations enriched for expression signatures associated with activated RAS signaling were selected. This is in keeping with prior work noting the association between RAS pathway mutations and monocytic differentiation ^18,19,42,43^, but also demonstrates that the association between RAS signaling activation and monocytic can be mutation independent. These observations are also consistent with the findings^24^ demonstrating selection for genetically distinct monocytic AML clones at relapse on a different venetoclax combination, Ven/hypomethylating agent (Ven/HMA). More surprisingly, we show that in our cohort, both immature and monocytic leukemia populations undergo further immunophenotypic and transcriptional shifts toward monocytic differentiation on Ven/Gilt treatment. This increase in monocytic differentiation appears tightly correlated with increased expression of activated RAS gene signatures, even in the absence of known new RAS pathway mutation, again suggesting that the underlying mechanism of Ven/Gilt resistance lies with the biologic consequences of increased RAS signaling.

In our *in vitro* model, both Ven-R and Ven/Gilt-R cell lines demonstrated an increase in RAS signaling and monocytic phenotype with resistance. For Ven/Gilt-R cells, an acquired S257L mutation in in RAF1, described in solid tumors but not hematologic malignancy^44^ likely drives increased MAPK signaling, and a subclonal BAX mutation, associated with venetoclax resistance in AML^37^, can also contributes to venetoclax resistance. However, no activating RAS pathway mutations or copy number alterations were identified in Ven-R cells. These data suggest that increase in RAS pathway activation and associated monocytic differentiation, whether associated with genetic mutations or not, can drive Ven resistance. Accordingly, we demonstrated in both *in vitro* models that inhibition of RAS signaling with the MEK inhibitor trametinib sensitized cells to both venetoclax and gilteritinib, a finding in line with previous observations that monocytic AML blasts resistant to venetoclax are sensitive to MEK inhibition^16,45^. Collectively, these data suggest that concurrent RAS pathway inhibition may effectively address resistance to venetoclax and Ven/Gilt.

Our findings also suggest that increased monocytic differentiation may occur independent of acquisition of new genetic drivers and that expression of monocytic gene expression programs occur without full expression of a monocytic immunophenotype. Our cell line data saw early upregulation of CD11b, an early to intermediate marker of monocytic differentiation, via both transcription and flow cytometry. This suggests that transcriptional signatures may be a more sensitive or early marker of monocytic differentiation and may explain why at least one study did not show a difference in response to venetoclax-based regimens based on clinical monocytic phenotype^46^.

At this time, it is unclear if increased expression of monocytic cell surface markers and monocytic differentiation programs associated with Ven/Gilt resistance serve as markers of increased RAS pathway activation or if monocytic differentiation state alone is sufficient to impart resistance. Monocytic immunophenotype has been linked to drug sensitivity in other contexts, both in association with sorafenib resistance^18^ and BET inhibitor sensitivity^47^. It is possible that RAS signaling activation associated with monocytic differentiation also underlies these observations. Regardless, the full molecular mechanisms underlying resistance to venetoclax mediated by either RAS or monocytic differentiation are not yet established. A previous study linked MCL1-dependent increased fatty acid metabolism to Ven/HMA resistance and RAS pathway mutations in AML LSCs^48^. Correspondingly, monocytic AML has also been linked to increased expression of MCL1 and MCL1-dependent oxidative phosphorylation gene signatures^17^. Additionally, monocytic LSCs have been shown to exhibit enriched MLL^17^ and purine metabolism gene signatures^24^.

In our Ven/Gilt-treated cohort, however, we were unable to demonstrate enrichment for oxidative phosphorylation or purine metabolism signatures in monocytic AML cells, and these signatures were also not enriched at relapse in either immature or monocytic AML cells. In contrast, we did observe that MLL signatures were enriched in monocytic compared to immature AML and in immature AML cells at relapse. Furthermore, in resistant cell line models of induced Ven versus Ven/Gilt resistance, although both models were characterized by increased RAS signaling and monocytic differentiation, we observed distinct transcriptional signatures. This included enrichment for an MLL signature in Ven-R, but not Ven/Gilt-R cells. These data suggest that, with the exception of activated RAS signaling, gene expression signatures previously linked to monocytic AML are not uniformly associated with resistance in all settings. This lack of uniformity suggest that these gene expression programs are not a cause of resistance to all venetoclax-containing combinations. As increasingly more venetoclax combinations are used, including expansion to triplet therapies including venetoclax, HMA, and a third targeted agent, characterizing these distinct resistance patterns becomes increasingly important.

Modulation of anti-apoptotic gene expression has been associated with RAS signaling activation and resistance to venetoclax. Specifically, it has been suggested that RAS/MAPK activation mediates resistance to both venetoclax and HMAs through MCL1 upregulation^12,49^. Loss of BCL2 and increased reliance on MCL1 to mediate survival have also been associated with monocytic differentiation and venetoclax resistance^17^. However, we found heterogeneous dynamics of MCL1 expression in our data, with increased expression in monocytic, but not immature, AML cells by SC analysis, and increased expression in Ven/Gilt-R relative to Ven-R cells. Upregulation of BCL2A1 has also been linked to resistance to both venetoclax^38^ and the FLT3 inhibitor quizartinib^50^. In our own data, we found increased BCL2A1 expression across heterogeneous subclones in Ven/Gilt resistant patient samples and in both Ven-R and Ven/Gilt-R cell lines. Similarly, independent overexpression of BCL2A1 in parental MOLM-14 conferred resistance to gilteritinib as well as the combination of venetoclax and gilteritinib. Together, these data offer BCL2A1 as a potentially promising target to combat resistance to Ven/Gilt and other targeted therapy combinations.

The specific role of monocytic cell surface markers in mediating Ven/Gilt resistance is unknown. We observed that upregulation of CD11b was associated with Ven/Gilt resistance across multiple heterogeneous subclones via both cell surface protein expression and transcription; CD11b was also upregulated in both of our resistance cell line models. CD11b, also known as Integrin alpha M (ITGAM), is a subunit of the heterodimeric integrin alpha-M beta-2 (α_M_β_2_), which has roles in both leukocyte migration and adhesion. Cell surface expression of CD11b/ITGAM mediates adhesion and spreading on α_M_β_2_ ligands^51^. Integrin alpha α_M_β_2_ is thought to have promiscuous interactions. In fact, CD11b has more than 100 reported ligands, including soluble mediators, counterreceptors, extracellular matrix (ECM) components^52^. In keeping with a potential role for CD11b in AML cell adhesion, patients with extramedullary leukemia are more likely to have blasts that express CD11b. Levels of CD11b expression have been linked to poor prognosis in AML^53^. Clinically, therapies that mobilize leukemic cells are thought to be sensitizing to chemotherapy, but the potential role of CD11b-mediated adhesion in modulating response to targeted therapies, such as Ven/Gilt, remains to be established.

Overall, our data emphasizes that AML cell state is dynamic. Differentiation has long been a marker of response to targeted therapies for certain AML subtypes, including acute promyelocytic leukemia^54^, and has more recently been observed with targeted IDH and menin inhibitors as well^55,56^. FLT3 inhibitors may induce differentiation^57^, though it may only be in a subset of cells^58^. For some targeted therapies, like venetoclax-based combinations, AML differentiation may instead be a marker of resistance. The idea of differentiation driving resistance has been long appreciated in solid tumors, including prostate and lung cancers^59,60^. Emerging data suggests that therapy-induced changes in cell state may also play an important role in AML. In pediatric AML treated with multi-agent cytotoxic chemotherapy, leukemias assume a more immature differentiation state at time of relapse^61^. Magrolimab-based regimens show erythroid differentiation at the time of resistance^62^. In this context, our data showing monocytic differentiation with targeted therapy suggests that while differentiation may be a widely applicable mechanism of resistance, the type of differentiation may vary with distinct therapeutic agents and combinations. In our secondary analysis of the BEAT AML cohort, we saw diverse changes in transcriptionally-defined cell state with treatment. Patients treated with a heterogenous array of targeted therapies showed decrease in progenitor-like and increase in monocytic-like gene programming. Looking forward, these dynamic cell state changes should be assessed prospectively to better understand the role in disease resistance or response.

While differentiation-based AML classification, like the French-American-British (FAB) schema, was described nearly half a century ago^63^, modern AML risk stratification is based on genetics^64^. This shift was based on data from patients treated with cytotoxic chemotherapy. In the era of targeted therapies, response prediction based on genetics alone may be insufficient, and a different risk stratification protocol based on a broader understanding of genetic and non-genetic resistance mechanisms may be warranted. This is particularly relevant in R/R disease, where treatment decisions come after one or more prior therapies, which may have selected for more diverse resistance mechanisms. Our data suggests that transcriptional differentiation state, at least in targeted therapy and venetoclax-containing regimens, may serve as a predictive biomarker of response.

Limitations of our study include that our DNA panel, while broad, was limited to hotspot mutations and our antibody panel did not utilize the full spectrum of monocytic markers, making it possible that some genetic mediators of monocytic immunophenotype and resistance could have been missed. For example, our SC DNA panel did not include the apoptosis effector BAX, for which inactivating missense or frameshift/nonsense mutations have been associated with venetoclax resistance in AML^37^. Further, we were unable to directly link mutational status with gene expression. Due to technical limitations of our CITE-seq analysis, we could not determine the distribution of RAS mutant cells within the leukemia populations with increased RAS signaling. Regardless, our findings add to a growing body of literature that suggests that RAS signaling activation is a central mediator of resistance to a wide swath of AML targeted therapies and that clinically effective RAS pathway inhibitors have potential to extend response for a large proportion of AML patients treated with these agents.

## Methods

### Patient samples

We studied 12 patients enrolled on the phase 1b trial of Ven/Gilt (NCT03625505), which included all trial patients with adequate serial samples available for analysis (**Figure S1**). From these 12 patients, there were 35 cryopreserved bone marrow or peripheral blood mononuclear cell samples. Details of the study design and outcomes have been previously published^9^. Study enrollment included informed consent for sample banking and analysis under protocols approved by each participating Institutional Review Board and conducted in accordance with the ethical standards of the institution as well as with the Declaration of Helsinki. All 35 samples were analyzed with SC DNA sequencing (DNA-seq) and 18 samples were analyzed with combined DNA and cell surface protein sequencing (DAb-Seq) (**Figure 1A**).

### Single Cell DNA and Protein Sample Preparation, Library Generation, and Sequencing

We performed SC DNA-seq or DAb-Seq on unsorted mononuclear cells from 12 patients across 35 samples using a microfluidic approach with molecular barcode technology using the Tapestri platform (MissionBio) as previously described^65,66^. Briefly, cryopreserved cells were thawed and normalized to 10,000 cells/μL in 180 μL PBS (Corning). Samples with more than one million viable cells recovered were additionally incubated with Human TruStain FcX (Biolegend) and salmon sperm DNA (Invitrogen) for 15 minutes at 4C. A pool of 9 oligo-conjugated antibodies (CD3, CD4, CD11b, CD13, CD19, CD33, CD34, CD117, CD45) (**Table S3**) was added, and cells were incubated for an additional 30 minutes. In addition, individual samples were also incubated with unique hashtags (BioLegend) to provide sample-level identifiers, and groups of 3 individual patients were pooled together in multiplexed runs.

Next, samples were resuspended in Cell Buffer (MissionBio), diluted to 4-7e6 cells/mL, and loaded onto a microfluidics cartridge, where individual cells were encapsulated, lysed, and barcoded using the Tapestri instrument. DNA from barcoded cells was amplified via PCR using a custom targeted panel that included 288 amplicons across 66 genes associated with acute leukemia generally and known clinical mutations in these samples specifically (**Table S2**). DNA PCR products were isolated, purified with AmpureXP beads (Beckman Coulter), used as a PCR template for library generation, and then repurified with AmpureXP beads. Protein PCR products (when applicable) were isolated from the supernatant from AmpureXP bead purification via incubation with a 5’ Biotin Oligo (ITD). Protein PCR products were then purified using Streptavidin C1 beads (Thermo Fisher Scientific), used as a PCR template for library generation, and then repurified using AmpureXP beads. Both DNA and protein libraries were quantified and assessed for quality via a Qubit fluorometer (Life Technologies) and Bioanalyzer (Agilent Technologies) prior to pooling for sequencing on an Illumina Novaseq.

### Single Cell DNA and Protein Data Processing

FASTQ files from single-cell DAb-seq samples were processed via an open-source pipeline as described previously^65^. This analysis pipeline trims adaptor sequences, demultiplexes DNA panel amplicons and antibody tags into single cells, and aligns panel reads to the hg19 reference genome. Valid cell barcodes were called using the inflection point of the cell-rank plot in addition to the requirement that 60% of DNA intervals were covered by at least eight reads. Variants were called using GATK (v 4.1.3.0) according to GATK best practices^67^. ITDseek was used to detect *FLT3* internal tandem duplication (ITD) from amplicon sequencing reads^68^. For each valid cell barcode, variants were filtered according to quality and sequence depth reported by GATK, with low quality variants and cells excluded based on the cutoffs of quality score < 30, read depth < 10, and alternate allele frequency < 20%. Cell-surface protein reads were normalized using centered log ratio transformations^69^.

### Single Cell DNA and DAbseq Downstream and Clonal Analysis

Prior to downstream analysis, pooled DAbseq runs were demultiplexed into individual patient samples using the SNACS algorithm, as previously described^70^. Briefly, this computational approach incorporates both sample-specific oligo-conjugated “hash” antibody tags as well as single nucleotide polymorphisms (SNPs) covered by the SC DNA panel to both demultiplex individual samples as well as identify and discard multiplets. Following de-multiplexing, for individual patients, we analyzed all variants present in >0.1% of cells. Variants were assessed for known or likely pathogenicity via ClinVar and COSMIC databases^71,72^ and previously identified, nonintronic somatic variants were included in clonal analyses. Genetic clones were defined as >10 cells possessing identical genotype calls, as per prior SC DNA studies^19,35,73^. Phylogenetic trees for individual patients were inferred using single cell inference of tumor evolution (SCITE), a probabilistic model for inferring phylogenetic trees^74^, using a global false positive rate set to 1% and a platform-provided false-negative rate as per prior SC DNA studies^35,75^. To define immunophenotypic subpopulations, unsupervised hierarchical clustering was performed using the scipy package in python, and UMAPs derived from protein expression data were constructed using the umap function with default settings.

### Single Cell CITEseq Sample Preparation

Cryopreserved bone marrow or peripheral blood mononuclear cells from 4 patients across 8 samples were thawed. For each specimen 1-2 million cells were blocked with Human TruStain FcX (BioLegend) and stained with a panel of TotalSeq-C antibodies from the Human Universal Cocktail (1 μg/each, BioLegend) in Cell Buffer (BioLegend) for 20 minutes at 4C. After staining, cells were washed in Staining Buffer, resuspended at 1,000 cells/μL, and immediately processed with the 10x Genomics controller and the Single Cell 5’ reagent kit with feature barcoding technology for cell-surface proteins. 5’ gene expression and feature barcoding libraries were constructed using a 10x Genomics library preparation kit. Library quality was assessed using Agilent High Sensitivity DNA kit and Bioanalyzer 2100, and libraries were quantified using dsDNA High-Sensitivity (HS) assay kit (Invitrogen) on Qubit fluorometer before being sequenced on an Illumina Novaseq. Targeted read depth for gene expression libraries was 100,000 reads/cell and for cell-surface libraries was 40,000 reads/cell.

### Single Cell CITEseq Processing and Annotation

Raw FASTQ data were aligned and filtered gene expression and feature barcode count matrices were generated using CellRanger (10x Genomics; v7.0.1) with the GRCh38 human reference genomic. Data filtering, integration, clustering, and differential expression analysis was performed using Seurat v5.0.0 in R^76^ Genes were filtered if detected in <3 cells and cells were filtered based on having low-complexity libraries (feature count < 200) or high mitochondrial content (>15%). Unsupervised cell clustering on transcriptional data was performed using Seurat with resolution set to 0.6, and clusters were visualized using the Seurat function RunUMAP with default settings.

Cell populations were annotated by a combination of clustifyr^77^, as per previously work annotating leukemic blast from SC RNAseq data^24^, and expression of cell-surface markers which can aid in distinguishing blasts from normal hematopoietic stem and progenitor cells^25–28^. Single cells classified as “unassigned”, “CD34+”, “CD14+”, or “CD16+” by clustifyr and expressing at least 2 of CD33, CD123, HLA-DR, or CLEC12A were considered leukemic blasts. To subclassify leukemic blasts into immature and monocytic populations, we next used six AML differentiation states first described by Van Galen et al by analyzing SC transcriptome data of multiple AML samples; importantly, this classification schema compared AML blasts against normal hematopoietic cells, thus affirming our classification of leukemic blasts vs normal cells^29^. For each single cell, we calculated a gene set score using the first principle component, as previously described^18,35^. Leukemic cells in the top 25th percentile for HSC- or Progenitor-like AML were classified as immature leukemia, while patients in the top 25th percentile for Monocyte-like AML were classified as monocytic leukemia. Differentially expressed genes for each cluster were determined using Seurat’s FindAllMarkers or FindMarkers functions, as appropriate.

### Gene Set Enrichment, CytoTRACE, and Pseudotime Analyses

Gene set enrichment analyses (GSEA)^78^ were performed using the fgsea R package (v1.15.1) with 1,000 gene set permutations by comparing single cells annotated as immature vs monocytic leukemia or by comparing pre-treatment vs relapse timepoints; all genes with log2FC ≥ ±threshold ≥ 0.1 were included. Gene sets used in this study included the molecular signatures database (MSigDB) hallmark v2022.1 (50 gene sets) and c2 (6,449 gene sets)^30,31^ as well as gene sets associated with specific cell differentiation states^29,32^ and venetoclax resistance^16^ in AML. Differentiation potential was determined using CytoTRACE v0.3.3, with 3,000 single cells sub-sampled from the 8 individual timepoints^33^. Pseudortime analysis was performed on all integrated cells using Lamian with default parameters^34^; Lamian was also used to perform repeated bootstrap sampling of cells along identified trajectories to calculate relative proportions at pre-Ven/Gilt and relapse timepoints.

### Cell Lines

Molm14 parental cells were a gift from Dr. Scott Kogan (University of California, San Francisco). Molm14 (*FLT3-ITD* mutant) and *NRAS* G12C co-mutant cells were generated by culturing Molm14 parental cells in media containing slowly escalating doses of quizartinib (0.5 - 20 nmol/L). Venetoclax/Gilteritinib (Ven/Gilt-R) and Venetoclax resistant (Ven-R) cells were generated by culturing quizartinib-selected *NRAS* G12C Molm14 cells in escalating doses of venetoclax (155 – 4454 nmol/L) with or without gilteritinib (14 – 34 nmol/L) (both compounds from Selleckchem)^79^. All resistant cells had confirmatory STR and Sanger sequencing performed.

Cell lines were cultured in RPMI-1640 (Gibco) with 10% fetal bovine serum (FBS) and 1% penicillin/streptomycin/L-glutamine (Gibco). Resistant cell lines were maintained in 4454 nmol/L venetoclax (ven-R cells) or venetoclax with 34 nmol/L gilteritinib (Ven/Gilt-R cells). Cells tested negative for *Mycoplasma* by the MycoAlert PLUS Mycoplasma Detection Kit (Lonza). Experiments were performed within three months of cell line thawing. Cell line authentication was performed at the University of California, Berkeley, DNA Sequencing Facility using short tandem repeat DNA profiling.

### Whole Exome Sequencing

Molm14 (*FLT3*-ITD mutant) and *NRAS* G12C co-mutant, Ven-R cells, and Ven/Gilt-R cells underwent whole exome sequencing to identify potential pathogenic mutations arising over the course of therapy selection. DNA was extracted from 1e6 cells from each cell line with the DNeasy Mini Kit (Quiagen). DNA was submitted to Novogene (Novogene Corporation Inc) for library preparation and sequenced on an NovaSeq X Plus sequencer (Illumina) sequencer. The resulting data was quality assessed using FastQC (v0.12.0), reads were trimmed with Trimmomatic (v0.38) to remove adapter and low average quality bases (<20) and sequences with minimum length below 50. Reads were aligned to the human reference genome (hg38) using BWA-MEM2 (v2.2.1). Picard (v2.12.3) was used to sort bam files and mark duplicates. GATK Mutect2 (v4.1.1) was used to call short somatic mutations (SNPs and indels) using Tumor with matched normal mode. Mutations were filtered using SNPsift (v5.2c) to remove low complexity regions (LCRs). Variants were annotated using SNPeff (v5.2c) and the databases dbSNP and ClinVar. Copy number variants were identified using CNVkit (v0.9.11).

### Bulk RNAseq Sample Preparation and Analysis

We performed bulk RNA-seq on Molm14/*NRAS* co-mutant non-resistant, Ven-R, and Ven/Gilt-R cells to identify transcriptional changes associated with acquired resistance. Total RNA was extracted from each triplicate (non-resistant and Ven-R) or quadruplicate (Ven/Gilt-R) sample with the RNeasy Mini Kit (Qiagen) and quality checked (RNA integrity number [RIN > 9). RNA was submitted to Novogene (Novogene Corporation Inc) for library preparation (NEBNext Ultra RNA Library Prep Kit for Illumina) and then sequenced in 100 bp paired-end read format on NovaSeq 6000 (Illumina, Inc.).

The resulting raw FASTQ data were then trimmed using Trimmomatic (v0.36) to remove adapter sequences and low-quality regions, mapped to a human reference genome (hg38) using the STAR alignment tool (v2.7.8a)^80^ and quantified using the featureCounts utility from the Subread package (v.1.5.1)^81^. Aligned reads were normalized using trimmed mean of M-values (TMM) from the EdgeR R package (v.3.24.3) and read counts were log2-transformed. The VOOM function from the limma package (v.3.38.3 was used to estimate the mean-variance relationship, assign precise weights, identify statistically significant differences in gene expression, and calculate statistical outputs. The fsgea package (v1.15.1) was used to perform pathway enrichment analysis.

### Cell Viability and Apoptosis Assays

For viability experiments, cells were seeded in three replicates in 96-well plates and exposed to increasing concentration of indicated drug (venetoclax or gilteritinib) +/-10 nM trametinib (Selleckchem) for 48 hours. Cell viability was assessed using CellTiter-Glo Luminescent Cell Viability Assay (Promega). Luminescence was measured on a Molecular Devices iD3 multimode plate reader and normalized to untreated controls.

For apoptosis experiments, cells were seeded in three replicates in 96-well plates and exposed to increasing concentration of indicated drug (venetoclax or gilteritinib) +/-10 nM of trametinib (Selleckchem) for 24 hours. CellEvent Caspase-3/7 Green Flow Cytometry Kit (#C10427, Thermo Fisher) was used to measure the ratio of apoptotic cells by flow cytometry (BD LSR Fortessa with HTS sampler). Percentage of caspase-3/7 negative cells in each treatment condition was normalized to vehicle-treated control populations (FlowJo v9, BD Biosciences).

### Flow Cytometry

To analyze cell surface phenotype, approximately 500,000 cells were stained with CD11b antibody (BioLegend, 301410) at 4oC for 30 minutes, washed with ice-cold FACS buffer, resuspended in FACS buffer, and analyzed on a NovoCyte 3005 flow cytometer. FCS files were analyzed on NovoExpress v1.6.1 (Agilent).

### Protein analysis

For immunoblotting, cultured cells were treated with indicated drug conditions for 24 hours, then lysed in buffer (50 mmol/L HEPES, pH 7.4, 10% glycerol, 150 mmol/L NaCl, 1% Triton X-100, 1 mmol/L EDTA, 1 mmol/L EGTA, and 1.5 mmol/L MgCl_2_) supplemented with protease and phosphatase inhibitors (#539134-1ML, 524625-1SET, EMD Millipore). Lysates were clarified by centrifugation, quantitated by BCA assay (#23225, Thermo Scientific), and normalized. 12 ug of protein per condition were run on 10% Bis-Tris gel (#WG1403A, Thermo Fisher) and transferred to nitrocellulose membranes (#162-0115, Bio-Rad). Actin was used as loading control.

Immunoblotting was performed using the following antibodies from Cell Signaling Technologies: anti-β-Actin (8H10D10, #3700), anti-Erk1/2 (3A7, #9107), anti-phoshpo-Erk1/2 (Thr202/Tyr204, #9101), anti-Akt (#9272), anti-phospho Akt (Thr308, #9275), anti-Stat5 (D2O6Y, #94205), anti-phospho-Stat5 (Tyr 694, #9351), anti-Bcl2 (124, #15071), anti-Bcl-xL (#2762), anti-Mcl-1 (#4572), anti-Bim (C34C5, #2933), anti-BMF (E5U2J, #50542). Cells then underwent incubation in secondary antibody (IRDye 800CW Goat anti-Rabbit, #935-32211, IRDye 680RD Goat anti-Mouse, #935-68070, LI-COR Biosciences) and imaging on an Odyssey CLx infrared imaging system (LI-COR Biosciences).

### Generation of BCL2A1 overexpressing cells

CRISPRa technology was used to create cell lines overexpressing BCL2A1. MOLM-14 CRISPRa cells were generated by transducing MOLM-14 parental cells with a lentiviral BFP-tagged VPR construct (JKNp44-pHR-SFFV-dCas9-HA-VPR-P2A-BFP, gift from Dr. Luke Gilbert, University of California San Francisco). Lenti-X 293T cells (Takara Bio) were infected using Lipofectamine 2000 (#11668019, Invitrogen) and viral supernatant harvested and filtered with 0.45 um nirocellulose membrane at 24 and 48 hours using Lenti-X concentrator (#631231, Takara Bio). Concentrated virus-containing media was used to spinfect MOLM-14 cells. After recovery, single clones were isolated using limiting dilution in a 96-well plate to establish cells with stable dCas9 VPR expression, confirmed by western blot. To create doxycycline-inducible expression vectors that were secondarily transduced into MOLM-14 CRISPRa cells, 24-bp oligonucleotides containing sgRNA sequences were synthesized (Integrated DNA Technologies), including 4bp overhangs (forward TCCC, complementary reverse AAAC), as previously described^82^. This enabled cloning into the Bsmb-I site of the lentiviral FgH1tUTG vector (Addgene #70183) using restriction digest and ligation of annealed oligonucleotides (protospacer sequences BCL2A1 g1: GCCTACGCACGAAAGTGACT, BCL2A1 g2: GGACACATGATGATACATGG). BCL2A1 overexpression was then confirmed via RT-PCR. Cells were incubated with or without doxycycline 1.0 ug/mL for 48 hours, and total RNA was isolated using RNeasy Mini kit (#74106, Qiagen). cDNA was synthesized from 2000 ng total RNA using SuperScript III reverse transcriptase (#18080044, Invitrogen) and qPCR was performed in 384-well plates in three technical replicates using Taqman Gene Expression assays on a QuantStudio 6 (Thermo FIsher). Differential gene expression was calculated using the 2^ΔΔCt^ method with GAPDH as the housekeeping control gene.

### Statistics and Reproducibility

Continuous variables were compared using Student’s *t*-test, paired *t-*test, or Wilcoxon ranked sums test and categorical variables were compared using chi-squared or Fisher’s exact tests. To evaluate clone-level cooccurrence, a contingency table was constructed for each mutation pair and the log2-transformed odds ratio computed; Fisher’s exact test was used to evaluate statistical significance. The association between individual mutations and cell-surface antibody expression was determined using point-biserial correlations. The association between gene set scores and between gene set scores and individual gene expression was determined using Spearman’s correlation, both in single-cell analysis and in bulk analysis of the BEAT AML data. All p-values for single-cell level comparisons were adjusted via the Bonferroni methods unless otherwise specified. All statistical analyses were performed in R (v. 4.0.2).

### Data Availability

The data discussed in this publication are in the process of being submitted to NCBI’s Gene Expression Omnibus; series accession number pending.

