## Supplemental Figures for "RAS pathway activation drives clonal selection and monocytic differentiation in FLT3 and BCL2 inhibitor resistance"

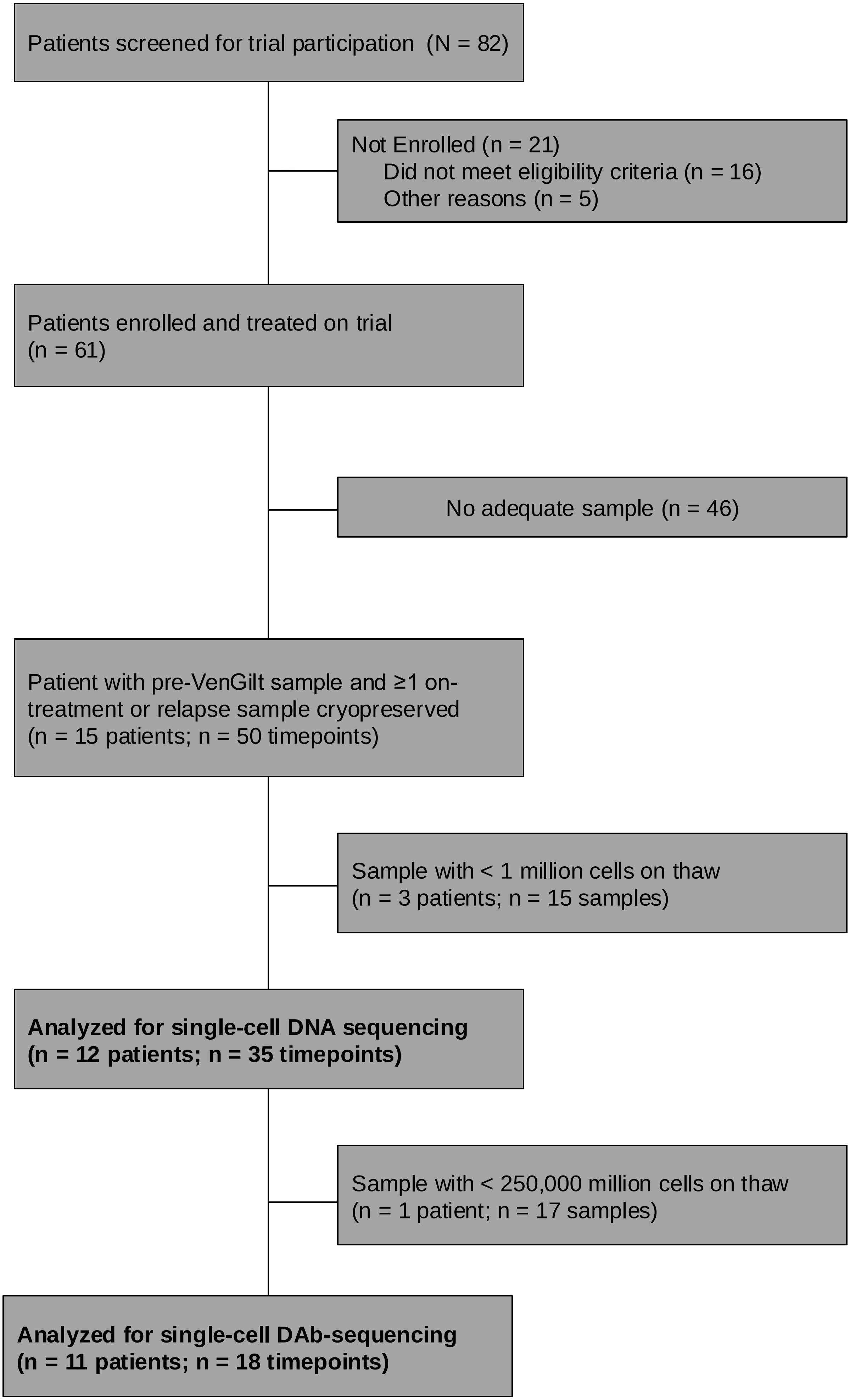


**Supplementary Figure 1.** Consort Diagram depicting patients enrolled on trial and with samples available for use in correlative single-cell analysis

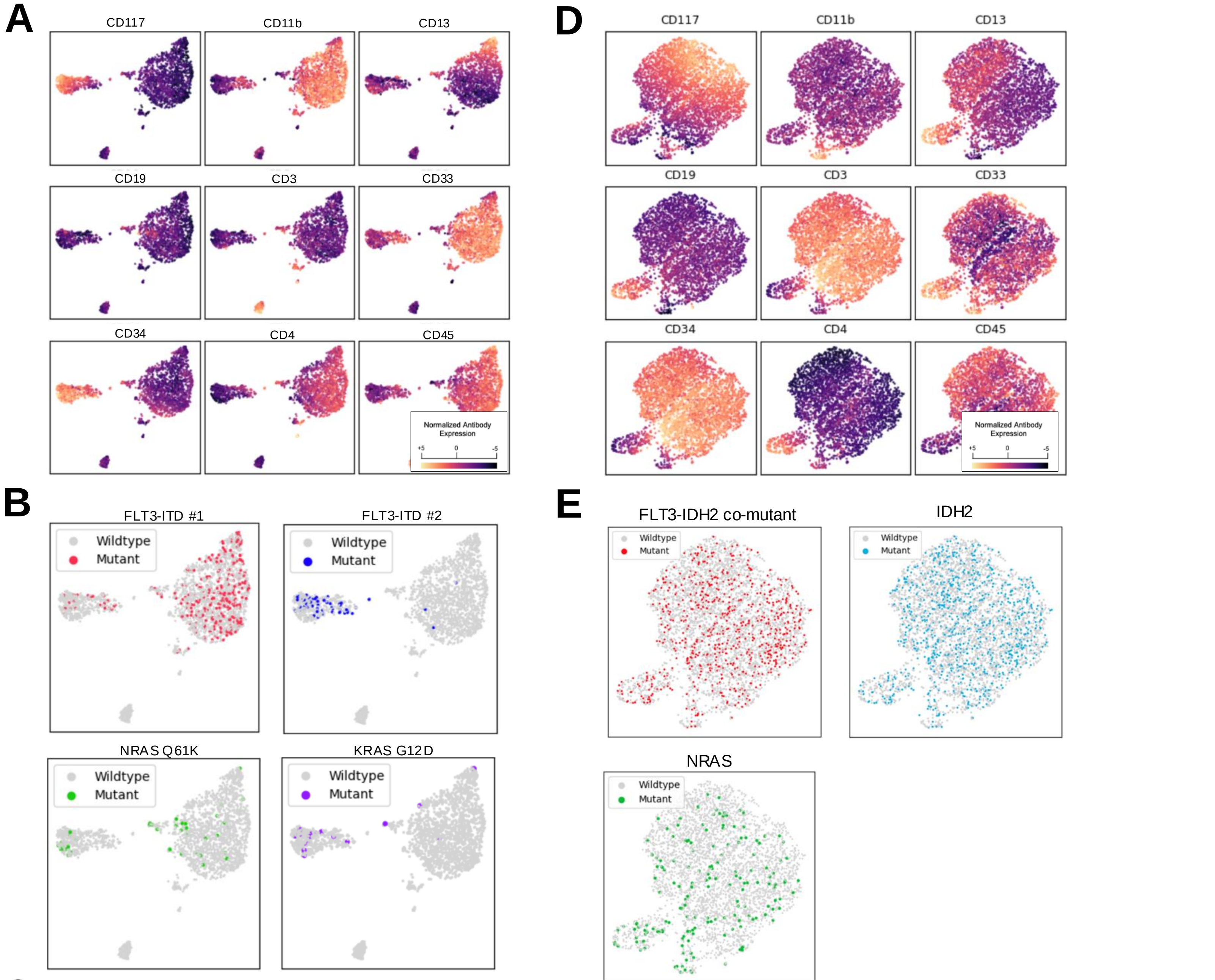


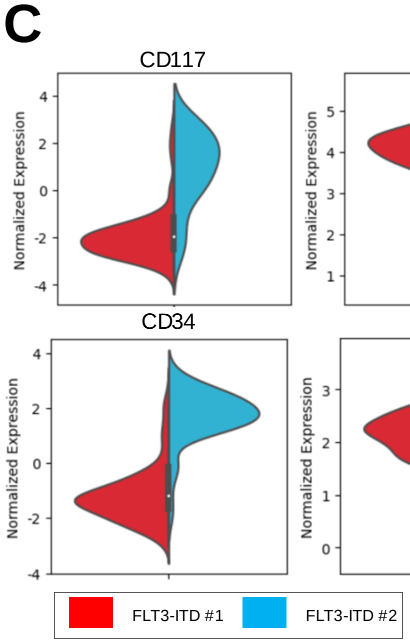


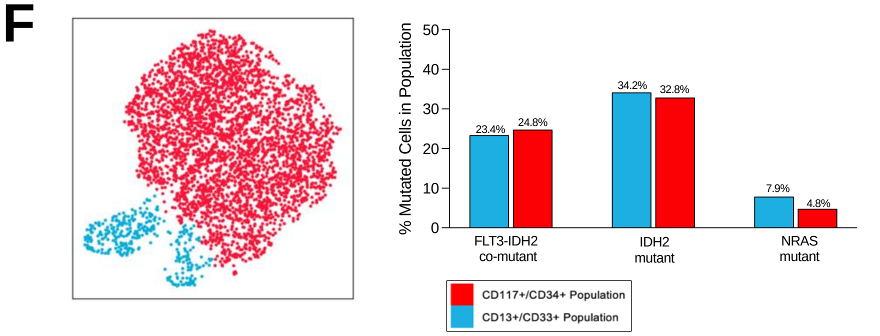


**Supplementary Figure 2. Diverse Genotype-Immunophenotype Associations**
***Left (Panels A – C):*** **Different genotypically-defined subclones demonstrated distinct immunophenotypes, including between 2 distinct FLT3 ITDs.** A. Immunophenotype-derived UMAP from 4,310 cells from pre-VenGilt sample from Patient 3. Cells are color-coded based on expression of cell-surface antibodies. **B.** UMAP from Panel A. Cells are color-coded based on genotype as FLT3-ITD#1 (red), FLT3-ITD #2 (blue), or NRAS (green). **C.** Split violin plot comparing cell-surface protein expression between 2 different *FLT3-ITD*-mutated subclones. ***Right (Panels D – F):*** **Different genotypically-defined subclones do not demonstrated distinct immunophenotypes, even when immunophenotypic heterogeneity is present.** **D.** Immunophenotype-derived UMAP from 3,825 cells from pre-VenGilt sample from Patient 2. Cells are color-coded based on expression of cell-surface antibodies. **E.** UMAP from Panel A. Cells are color-coded based on genotype as FLT3-ITD/IDH2 co-mutated (red), IDH2 (blue), or NRAS (green). **F**. *Left:* Hierarchical clustering of immunophenotype-derived UMAP defines 2 distinct populations, a CD117+/CD34+ population and a CD13+/CD33+ population. *Right:* For all the FLT3/IDH2 and IDH2 mutant clones, there is no significant difference in proportion of mutated cells within each immunophenotypic population. NRAS-mutated clones are enriched in the CD13+/CD33+ population.

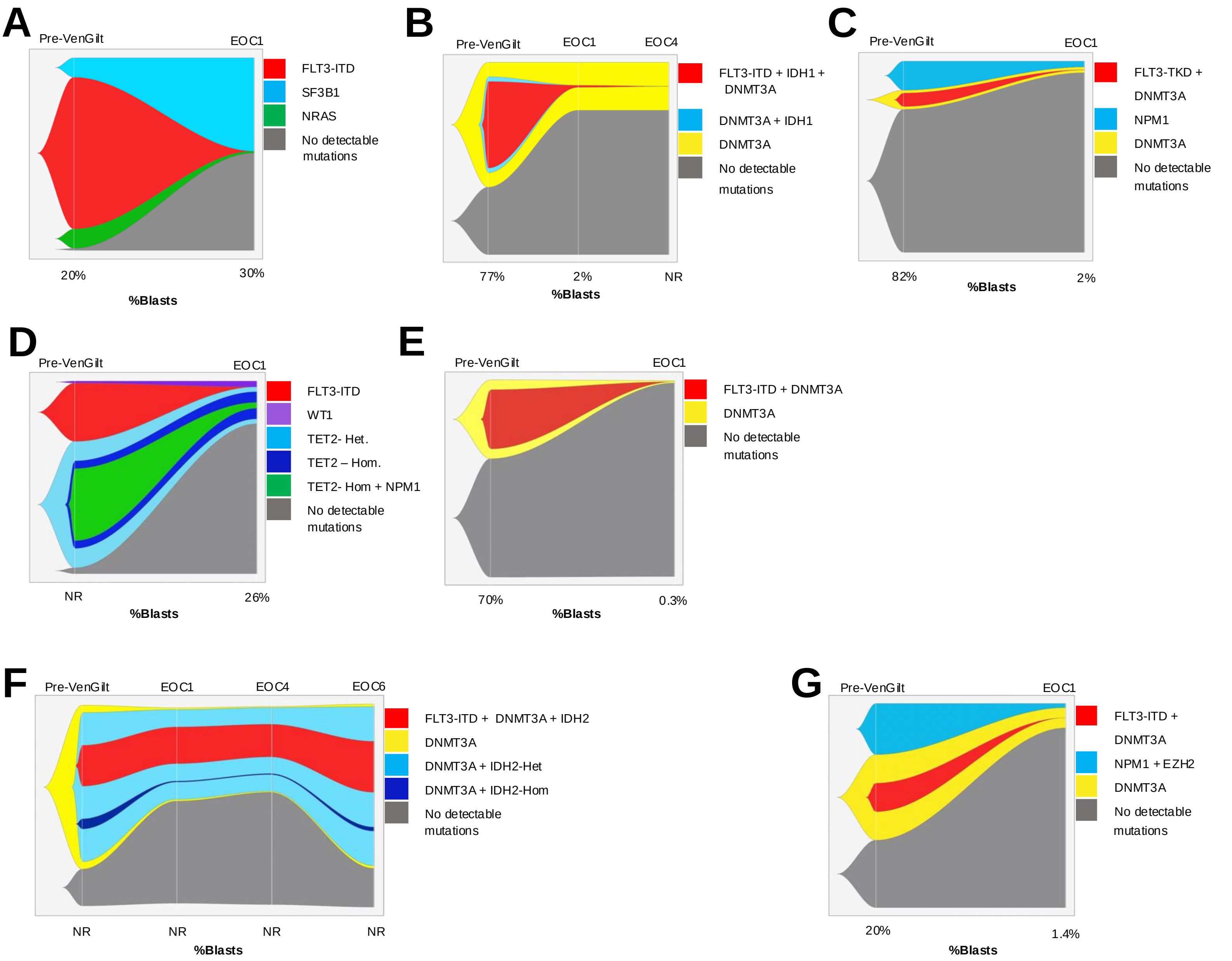


**Supplementary Figure 3. Genetic Clonal Evolution for 7 patients treated with Ven/Gilt therapy on clinical trial.** Remaining 5 patients are in Figure 2C-G. Fishplots illustrating dynamic clonal architecture as measured by SC analysis. For each plot, top x-axis indicates clinical timepoint (EOC, end of cycle) and bottom x-axis indicates percent blasts on corresponding clinical bone marrow biopsy, if available (NR, Not Reported).
***Patients in which FLT3-mutated clones become undetectable:*** **A.** 7,192 SCs from Patient 7 **B.** 3,222 SCs from Patient 8 **C.** 5,261 SCs from Patient 9 **D.** 632 SCs from Patient 10 **E.** 2,309 SCs from Patient 12. See also Figure 2D, G.
***Patients in which FLT3-mutated clones decrease in size but do not become undetectable***: **F.** 4,912 SCs from Patient 6 **G.** 16,546 SCs from Patient 11. See also Figure 2F.
***Patients in which one FLT3-mutated clone becomes undetectable and one does not:*** See Figure 2C, E.

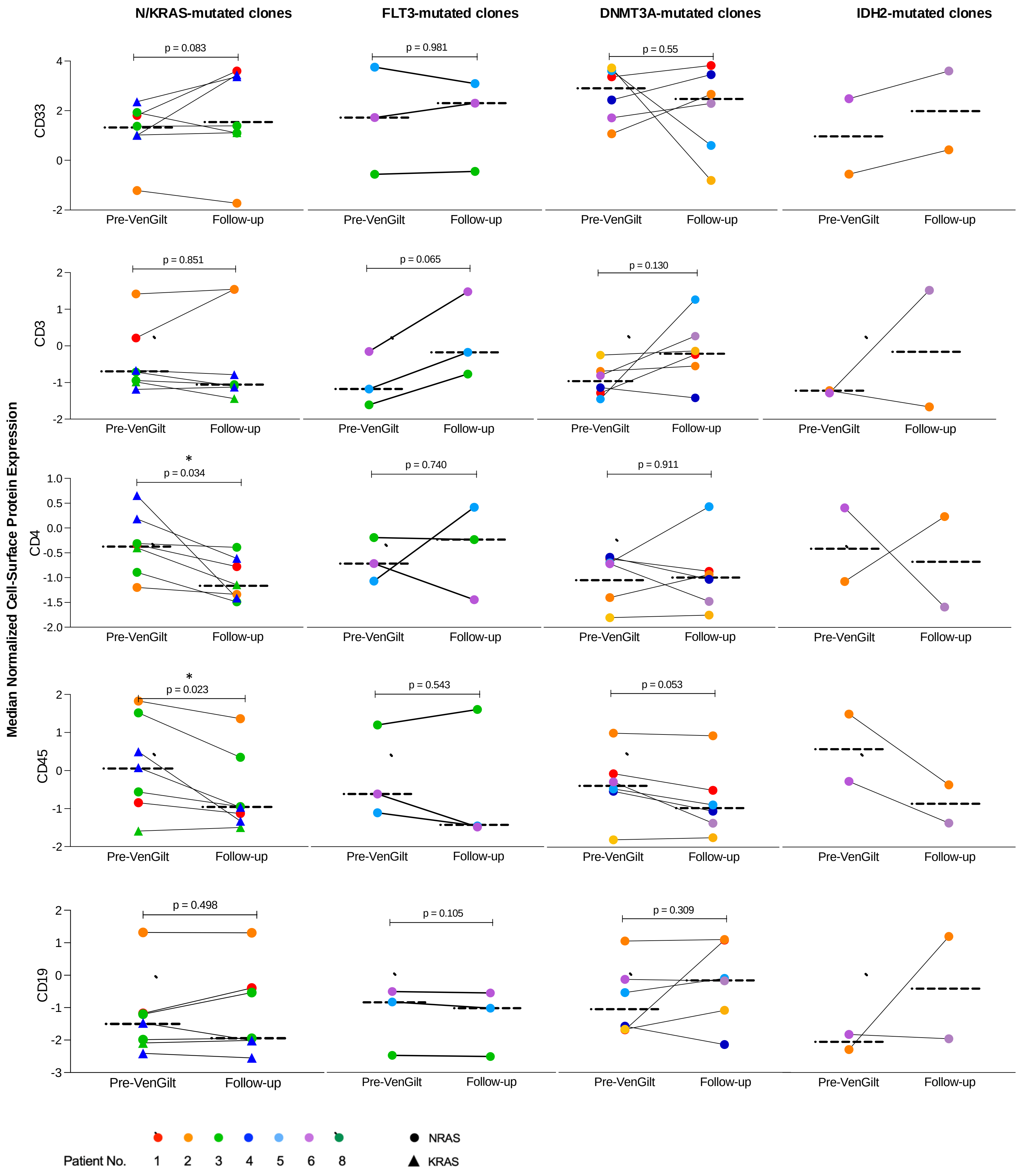


**Supplementary Figure 4.** Dot plot comparing pre-Ven/Gilt and post-Ven/Gilt samples from patients with *NRAS, KRAS, FLT3, DNMT3A,* or *IDH2*-mutated clones. The y-axis is median normalized expression of CD33, CD3, CD4, CD45, and CD19 cell-surface proteins. Dashed lines indicate medians and dots are color and shape-coded based on source patient and/or mutation type. Significance from paired T tests indicated as *p < 0.05.

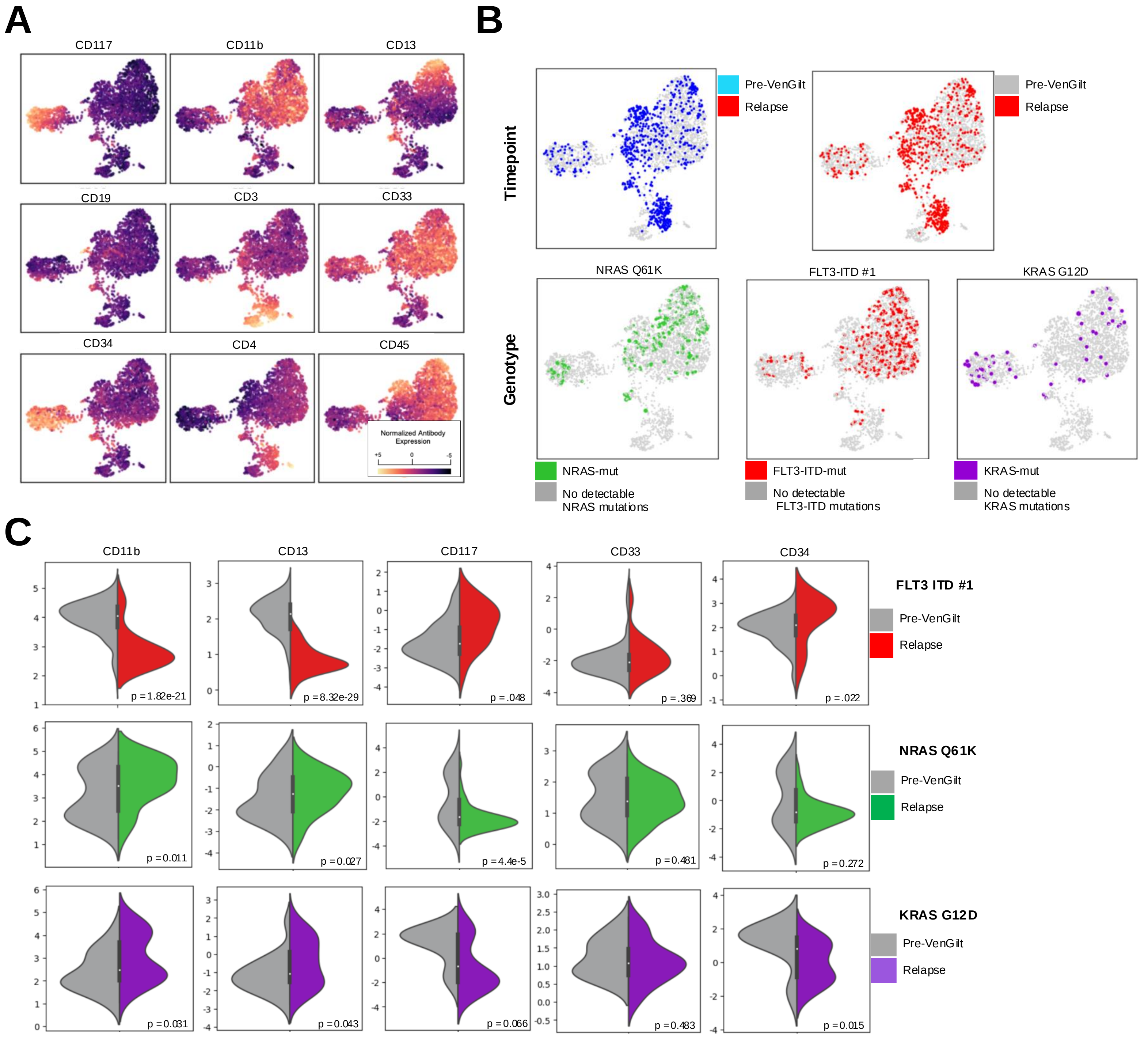


**Supplementary Figure 5. RAS mutated, but not FLT3-mutated, clones demonstrate increased expression of monocytic markers with therapeutic pressure.** In Patient 3, detailed in this figure, resistant *NRAS* and *KRAS* clones both demonstrated upregulation of CD11b and CD13 with therapeutic pressure; furthermore, the *NRAS*-mutated clone demonstrated decreased expression of immature marker CD117 and the *KRAS*-mutated clone demonstrated decreased expression of the immature marker CD34. This pattern was not observed in the *FLT3* mutated clone from the same patient, where CD11b and CD13 increased while CD34 decreased.

**A.** Immunophenotype-derived UMAP from 5,606 cells from integrated pre- and post-VenGilt samples from Patient 3. Cells are color-coded based on expression of cell-surface antibodies

**B**. UMAP from Panel A. Cells are color coded based on timepoint (top row) and genotype (bottom row).

**C.** Split violin plot comparing expression of select cell-surface proteins (CD11b, CD13, CD117, CD33, CD34) for pre-VenGilt vs post-VenGilt timepoints for 3 genetically defined subclones: FLT3-ITD#1 (top row), NRAS Q61K (middle row) and KRAS G12D (bottom row). While all clones demonstrated decreased CD117 and CD34 expression, only NRAS and KRAS mutated clones demonstrated increased CD11b and CD13 expression.

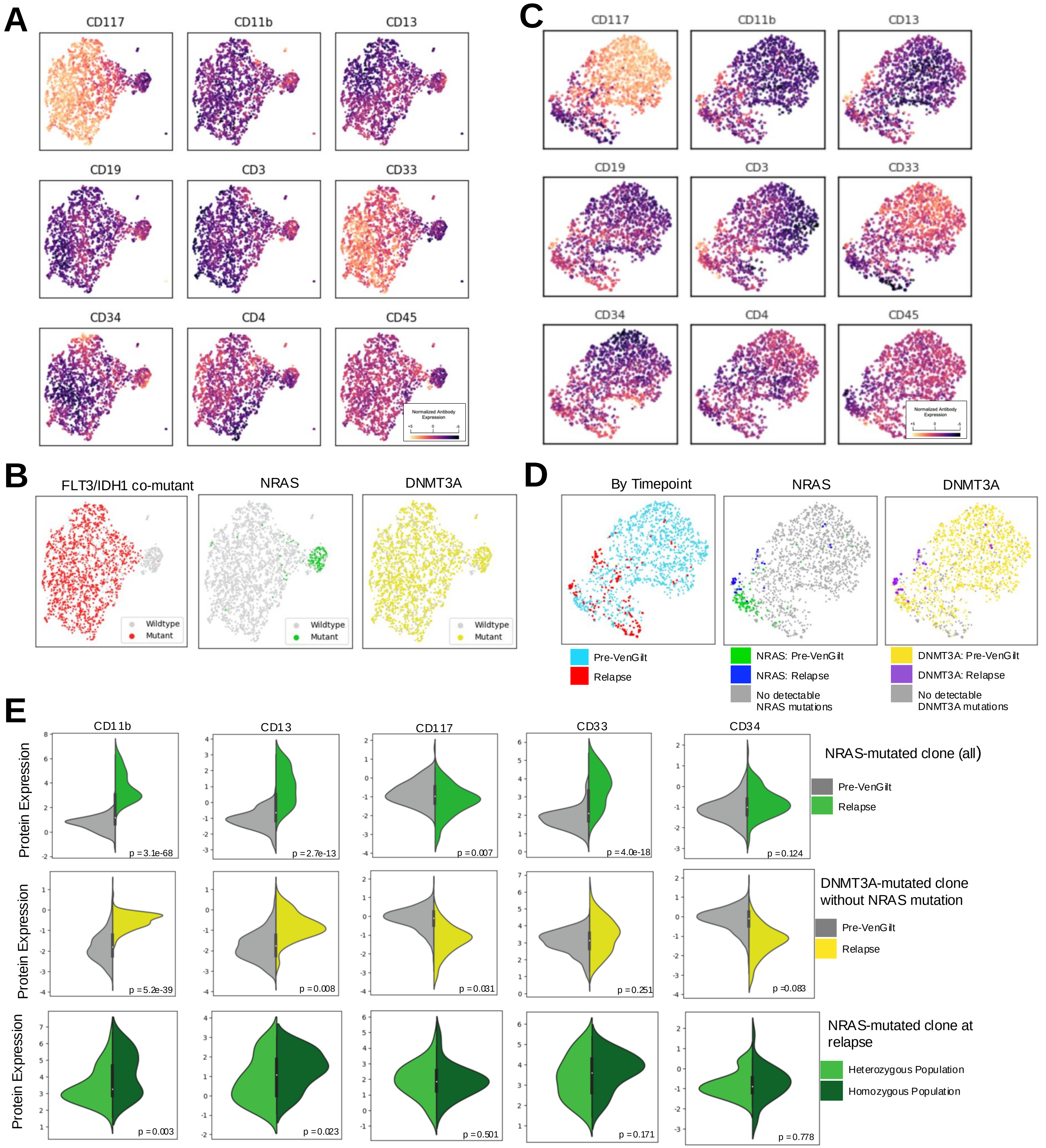


**Supplementary Figure 6. NRAS is associated with monocytic markers at baseline; NRAS and non-NRAS mutant cells become more monocytic with therapeutic pressure.** In Patient 1, detailed in this figure, a *NRAS*-heterozygous population is associated with a monocytic immunophenotype pre-Ven/Gilt. With treatment pressure, all *NRAS*-mutated cells developed increased expression of CD11b and CD13; this increased expression was even high in the *NRAS*-homozygous population. Interestingly, in this patient, *DNMT3A*-mutated cells without detectable co-mutations also demonstrated increased CD11b and CD13 expression with corresponding decreased in the the expression of immature marker CD117.

**A.** Immunophenotype-derived UMAP from 2,913 cells from pre-VenGilt sample from Patient 1. Cells are color-coded based on expression of cell-surface antibodies. **B**. UMAP from Panel A. Cells are color-coded based on genotype as FLT3-IDH1 co-mutant (red), NRAS (green) or DNMT3A (yellow). **C.** Immunophenotype-derived UMAP of 5,817 cells integrated from pre-VenGilt and relapse samples from Patient 1. Cells are color-coded based on expression of cell-surface antibodies. **D.** UMAP from Panel C. Cells are color-coded based on timepoint (left), NRAS genotype plus timepoint (center), and DNMT3A genotype plus timepoint (right). **E.** Split violin plot comparing expression of CD11b, CD13, CD117, CD33, and CD34 for cells. The top top compares protein expression for the NRAS-mutated clone pre-VenGilt (grey) vs relapse (green), the center row compares protein expression of the DNMT3A-mutated clone without co-mutations pre-VenGilt (grey) vs relapse (yellow), and the bottom row compares expression of the heterozygous (light green) vs homozygous (dark green) NRAS-mutated clone at relapse.

A

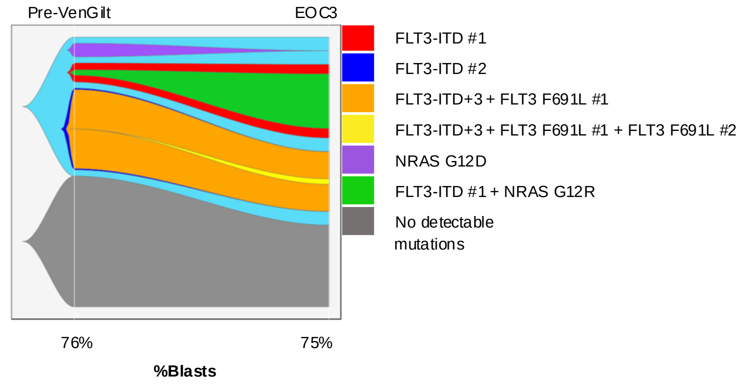


B

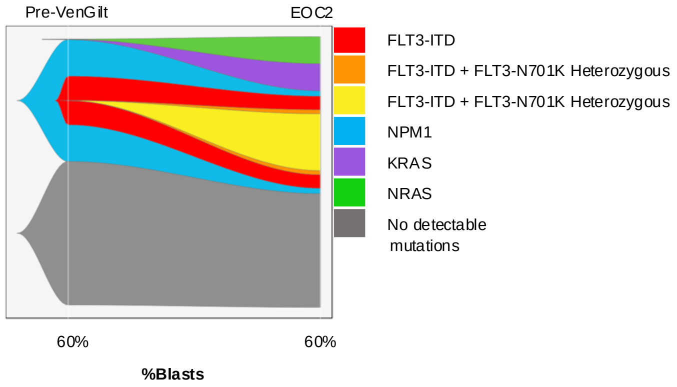


**Supplementary Figure 7. Genetic Clonal Evolution for 2 patients treated with Ven/Gilt therapy not on clinical trial.** Fishplots illustrating dynamic clonal architecture as measured by SC DNA analysis. For each plot, top x-axis indicates clinical timepoint (EOC, end of cycle) and bottom x-axis indicates percent blasts on corresponding clinical bone marrow biopsy. **A.** 2,413 SCs sequenced from Patient 13. **B.** 1,730 SCs sequenced from Patient 14. A version of the fishplot from Patient 14 has been previously published[82]

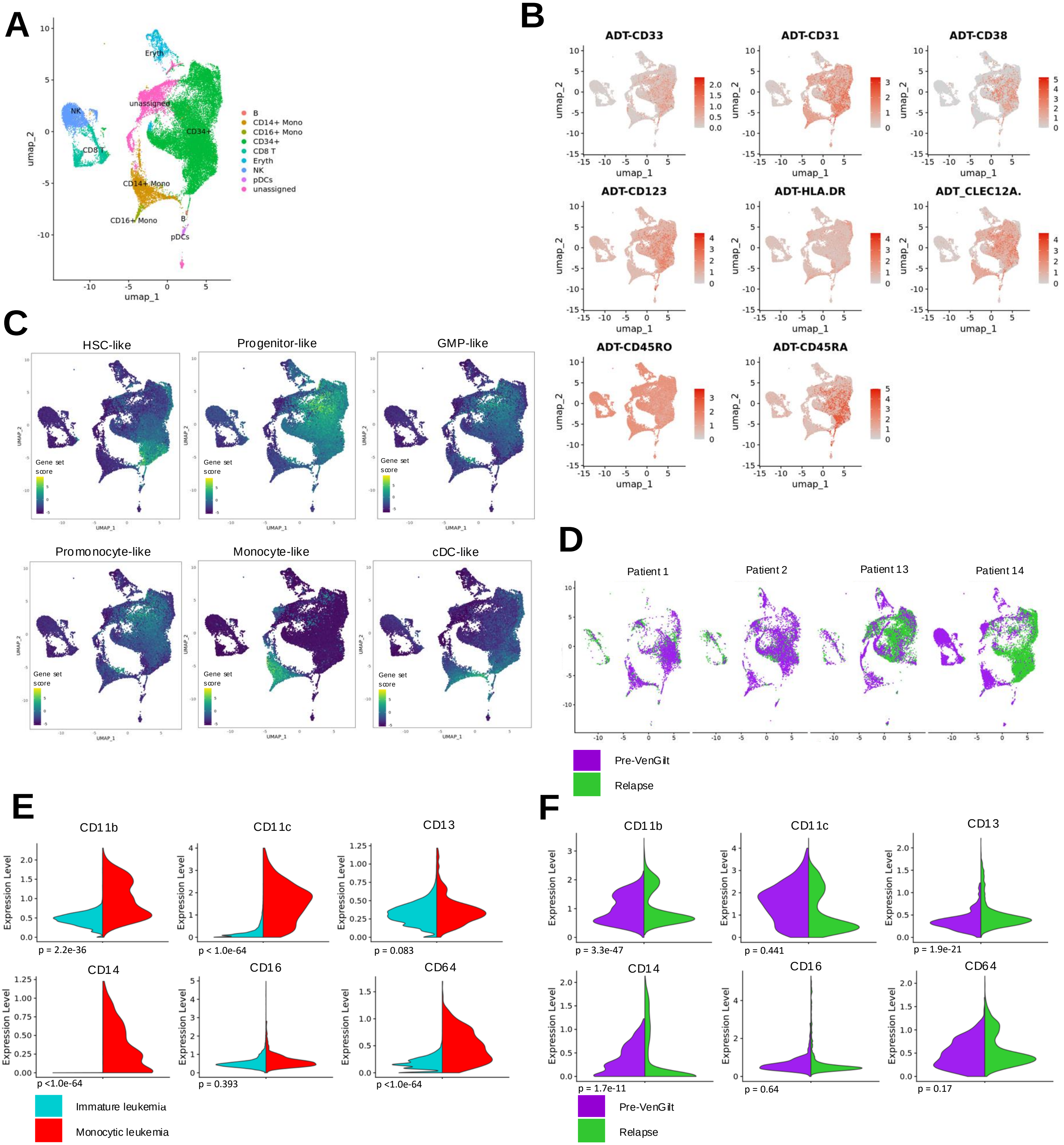


**Supplementary Figure 8. Classification of integrated SC CITEseq data identifies leukemic blasts and blasts subsets, which demonstrate distinct and dynamic immunophenotypes.**
**A.** UMAP derived from transcriptional data from 42,853 single-cells from pre-treatment and relapse timepoints from 4 patients treated with VenGilt annotated with clustifyr framework**. B.** RNA-derived UMAP from Panel A. Cells are color-coded based on expression antibody-derived tags (ADTs) from red (greatest cell-surface protein expression) to grey (least). **C.** RNA-derived UMAP from Panel A. Cells are color-coded based on gene set score derived from gene expression signatures from 6 AML subtypes. **D.** UMAP from Panel A split by parent sample. Cells are color-coded based on timepoint. **E.** Split violin plot comparing normalized cell surface antibody expression of monocytic markers between the immature vs monocytic leukemia populations at baseline and prior to VenGilt therapy. **F.** Split violin plot comparing normalized cell surface antibody expression of monocytic markers between Pre-VenGilt vs Relapse timepoint for the monocytic leukemia population.

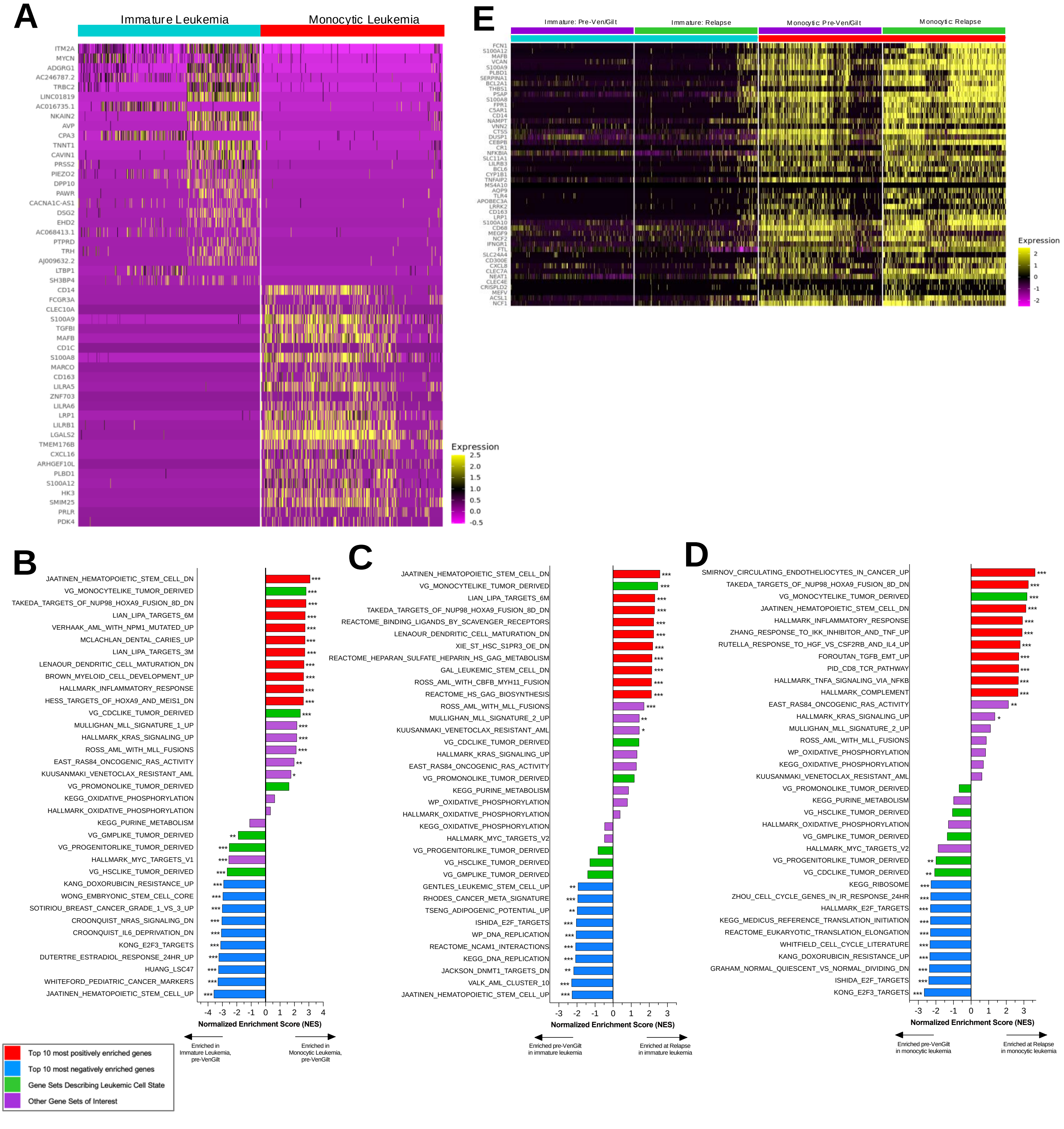


**Supplementary Figure 9. Immature and Monocytic Leukemia Populations demonstrate distinct transcriptional signatures, and demonstrate enrichment for monocyte-like AML with therapeutic resistance**

**A.** Heatmap of scaled expression values for the 25 genes with greatest upregulation in single cells from the immature (left columns) vs monocytic (right columns) leukemia populations.

**B.** Bar plot of normalized enrichment scores (NES) from Gene Set Expression Analysis (GSEA) of **all** leukemic cells. Positive enrichment indicates enrichment in the monocytic leukemia population and negative enrichment indicates enrichment in the immature leukemia population. **C.** Bar plot of normalized enrichment scores (NES) from Gene Set Expression Analysis (GSEA) of **immature** leukemia cells. Positive enrichment indicates enrichment at relapse and negative enrichment indicates enrichment pre-VenGilt. **D.** Bar plot of normalized enrichment scores (NES) from Gene Set Expression Analysis (GSEA) of **monocytic** leukemia cells. Positive enrichment indicates enrichment at relapse and negative enrichment indicates enrichment pre-VenGilt. For all bar plots, the top 10 positively enriched gene sets are color-coded in red, the top 10 negatively enriched in blue, gene sets derived from AML differentiation states are in geen, and additional gene sets of interest in purple. Statistical significance is indicated as ***q <0.001, ** q <0.01, *q <0.05

**E.** Heatmap of scaled expression values of 30 genes comprising the van Galen monocyte-like AML transcriptional signature for the immature leukemia population pre-VenGilt, the immature leukemia population at relapse, the monocytic leukemia population pre-VenGilt, and the monocytic leukemia population at relapse.

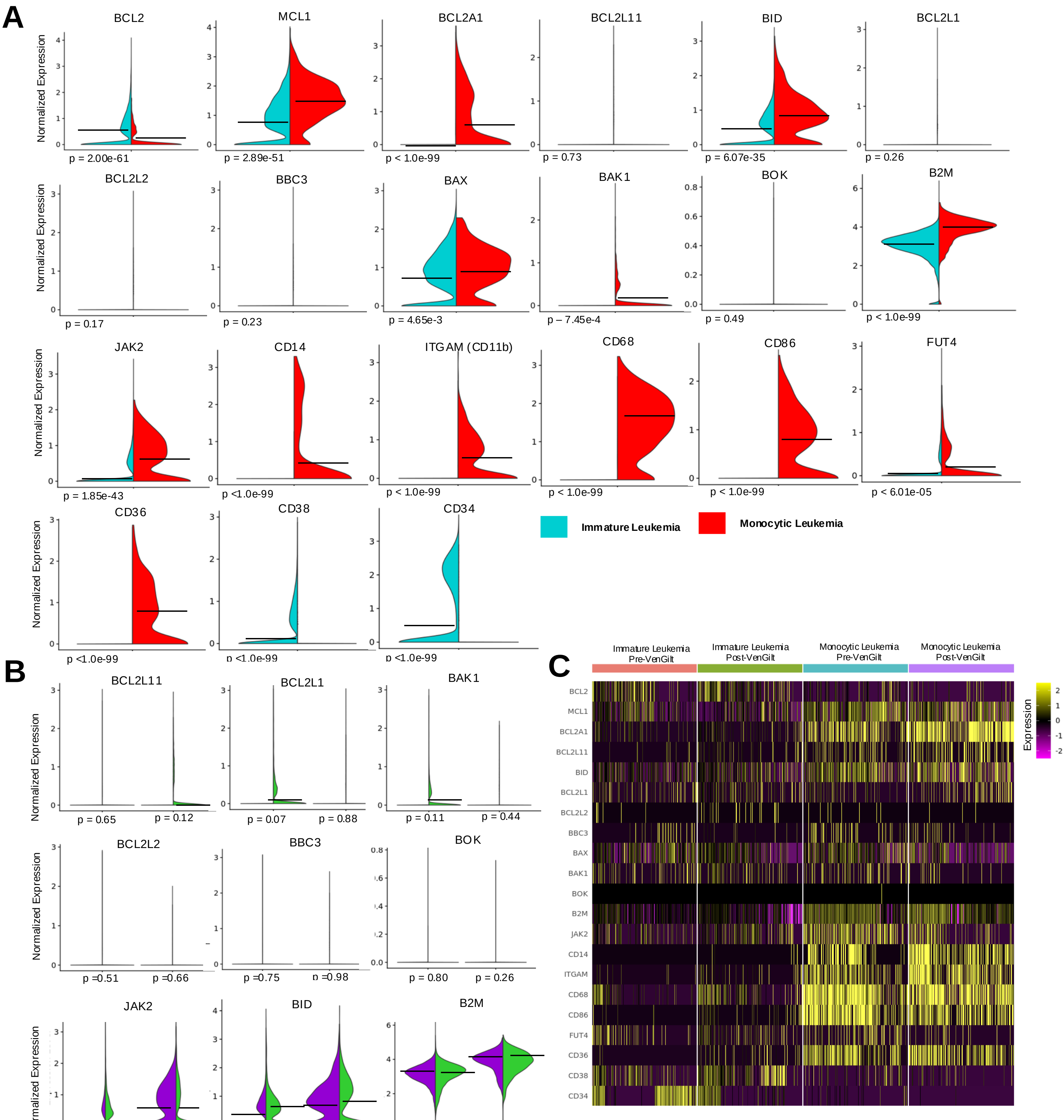

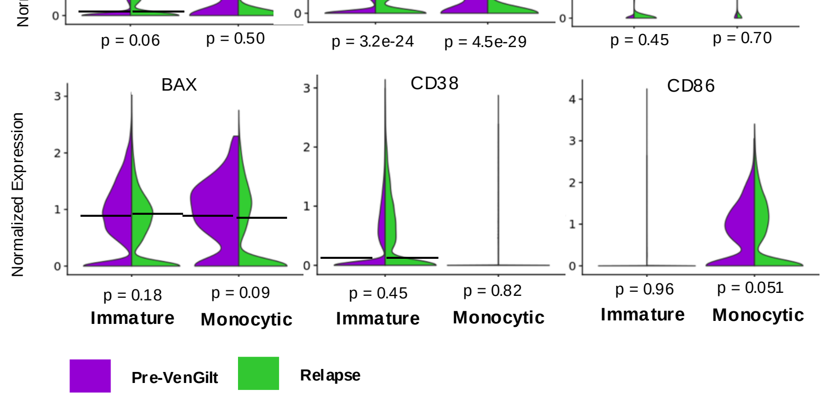


**Supplementary Figure 10. Genes associated with Venetoclax resistance are associated with monocytic cell state and demonstrate distinct expression patterns in immature vs monocytic populations with therapy resistance**

A. Split violin plot comparing expression of genes associated with Venetoclax resistance in the literature in the immature vs monocytic leukemia populations at baseline and prior to VenGilt therapy

B. Split violin plot comparing comparing expression of genes associated with Venetoclax resistance in the literature at baseline vs relapse in both immature and monocytic populations. Additional genes in Figure 5C.

C. Heatmap of scaled expression values of genes associated with Venetoclax resistance for the immature leukemia population pre-VenGilt, the immature leukemia population at relapse, the monocytic leukemia population pre-VenGilt, and the monocytic leukemia population at relapse.

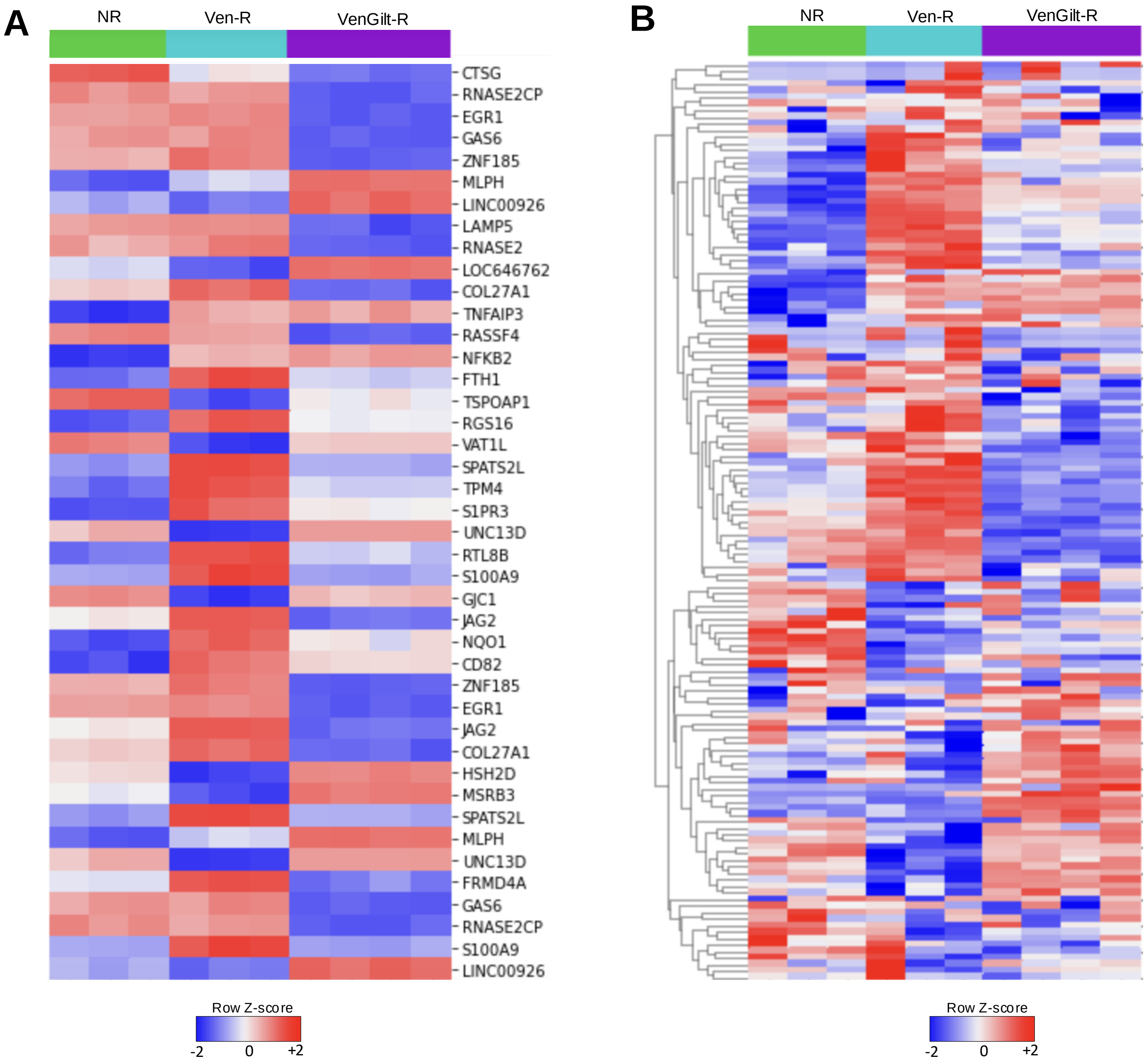


**Supplementary Figure 11. Generation of Venetoclax/Gilteritinib resistant (VenGilt-R) and Venetoclax resistant (Ven-R) *FLT3-ITD/NRAS G12C* co-mutant Molm14s demonstrates unique transcriptional profiles**

A. Heatmap of normalized gene expression of 50 most significant differentially expressed genes between VenGilt-R vs non-resistant (NR) cells, Ven-R vs non-resistant cells, and VenGilt-R vs Ven-R resistant cells.

B. Heatmap of normalized gene expression of all genes comprising the Hallmark KRAS UP gene signature for VenGilt-R, Ven-R, and non-resistant cells

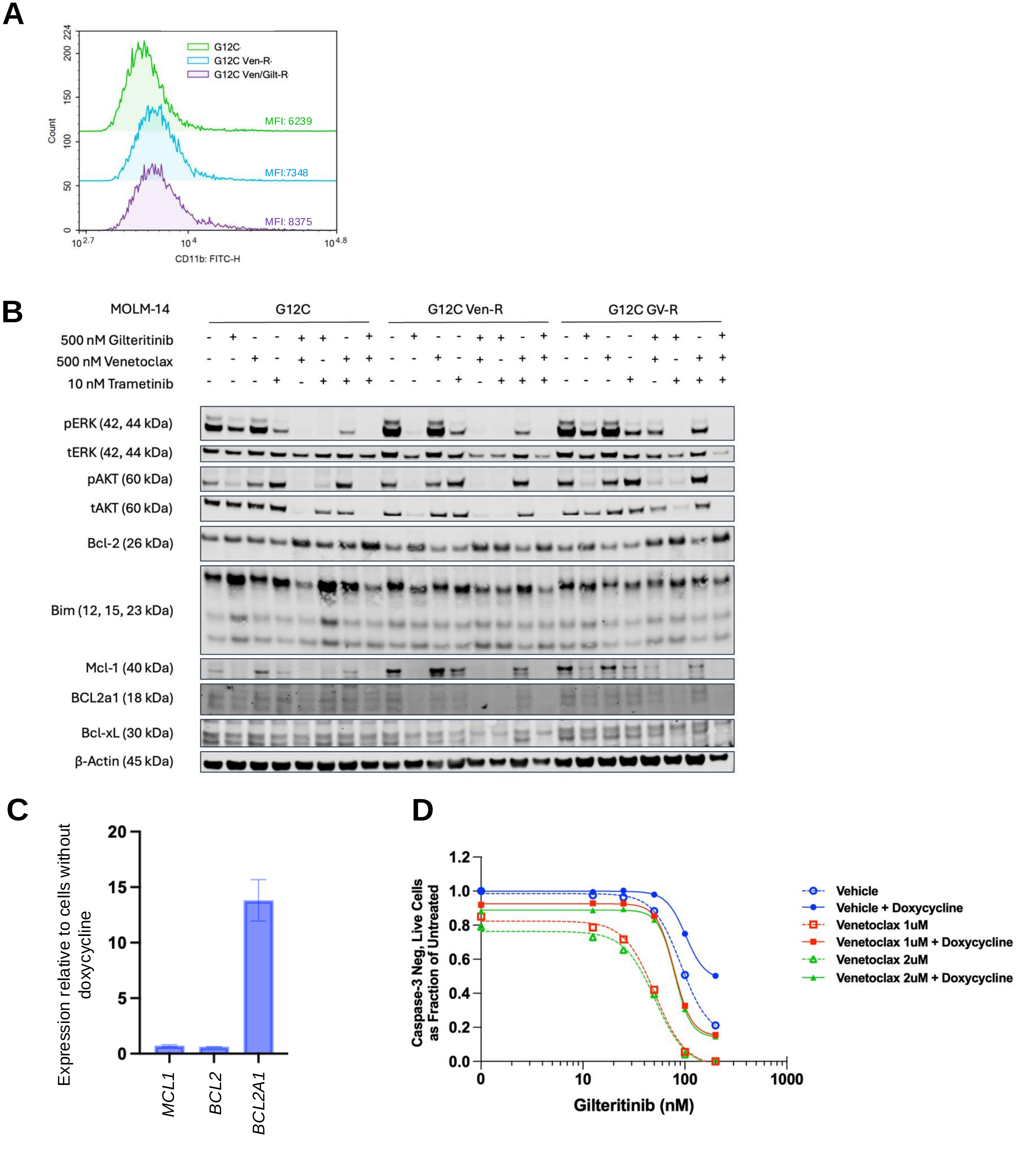


**Supplementary Figure 12. Expression of proteins associated with monocytic differentiation and/or venetoclax resistance in Venetoclax/Gilteritinib resistant (VenGilt-R) and Venetoclax resistant (Ven-R) *FLT3-ITD/NRAS G12C* co-mutant cell lines**

A. Cell surface flow cytometry of MOLM-14 (FLT3-ITD-mutant) cell lines with NRAS G12C co mutation demonstrates increase in CD11b (*left)* but not CD16 *(right)* cell surface markers in cells selected for resistance to Ven or VenGilt relative to unselected cells.
B. Western blot analysis of indicated proteins (phosphorylated ERK [pERK], total ERK [tERK], pAKT, tAKT, BCL2, BIM, MCL1, BCL2A1, BCLxL) in cell lines exposed to 500nM Gilt, 500 nM Ven, or 10nM trametinib as indicated
C. Expression of MCL1, BCL2, and BCL2A1 transcripts by RT-PCR in MOLM-14 cells engineered to express BCL2A1 in the presence of doxycycline. Expression levels are shown after 48 hours of exposure to 1uM/mL doxycycline.
D. Dose response curves representing caspase 3/7 expression of *BCL2A1* over-expressing parental MOLM-14 cells from Panel D with 0, 1, and 2 uM venetoclax and increasing doses of gilteritinib, with and without doxycycline induction.

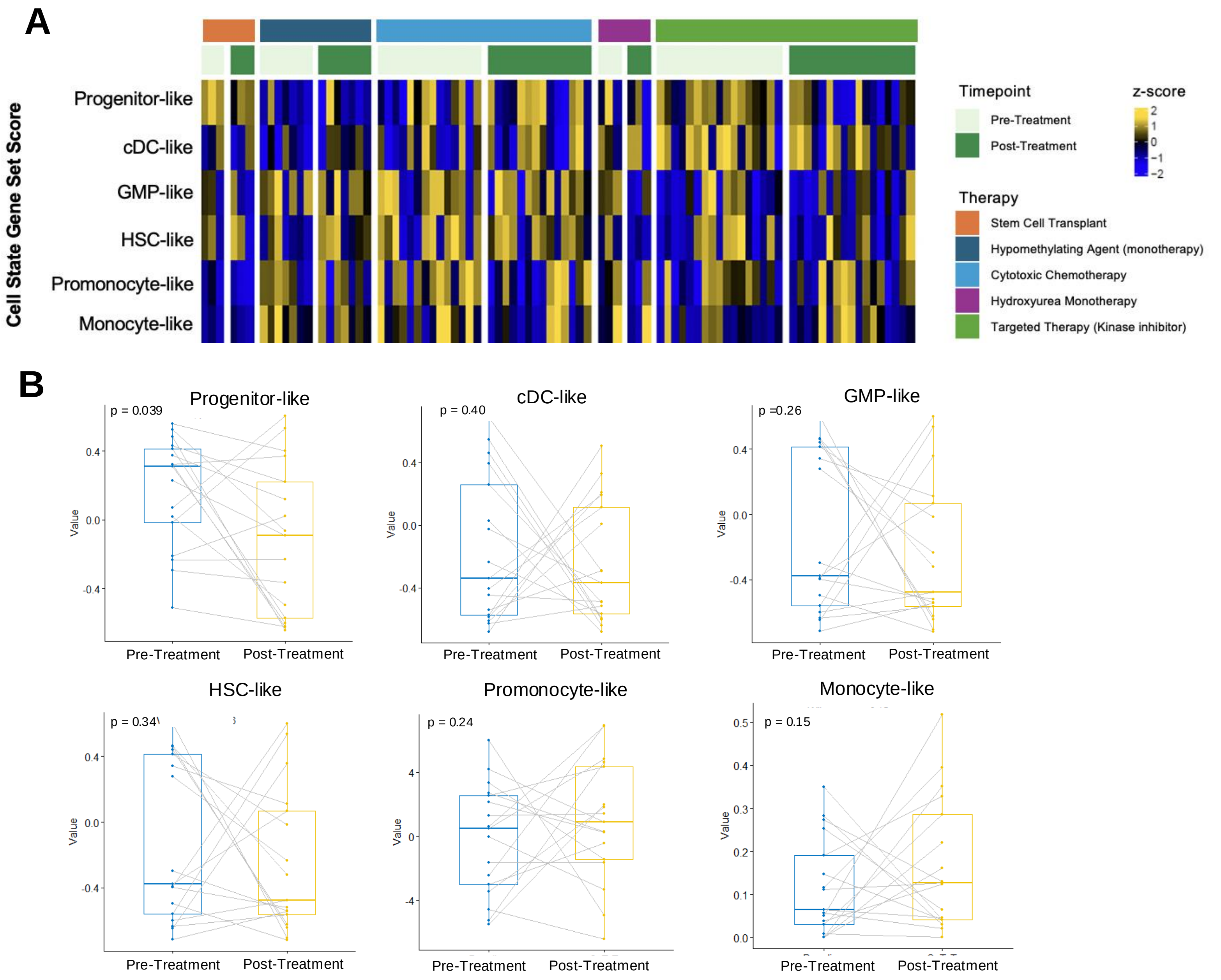


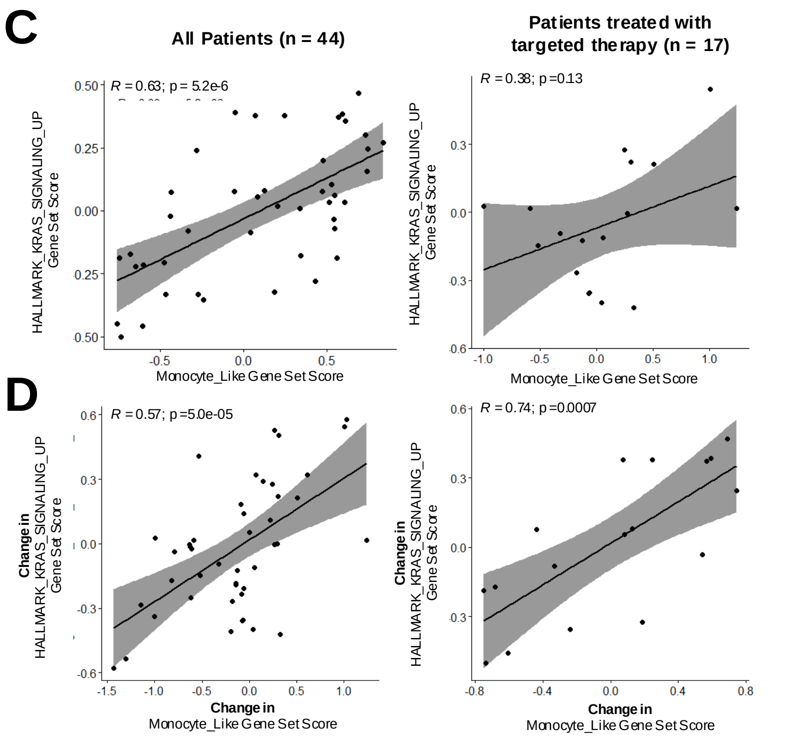


**Supplementary Figure 13. Dynamic cell state and monocytic differentiation are observed in the BEAT AML cohort despite heterogenous treatment**

**A.** Heatmap of normalized expression of gene set score derived from transcriptional cell state signatures across 44 patients with longitudinal samples from the BEAT AML cohort. Each column is a unique patient timepoint. Columns are grouped by treatment type and timepoint (pre-treatment and post-treatment).

**B.** Box plot of paired pre- and post-treatment gene set scores derived from transcriptional cell state signatures for 17 patients from the BEAT AML cohort treated with targeted therapies. With therapy, there was a significant decrease in progenitor-like cell state.

**C.** Scatterplot of gene set scores for the Hallmark KRAS signaling signature vs Monocyte-like gene signature at baseline for all patients (*left*) and for patients treated with targeted therapy (*right*).

**D.** Scatterplot of change in gene set scores for Hallmark KRAS signaling vs Monocyte-like gene signatures between post-treatment and pre-treatment timepoints for all patients (*left)* and patients treated with targeted therapy (*right*). With treatment pressure, and increase in KRAS signaling was tightly correlated with an increase in monocyte-like cell state.
